# Transforming the cytokine literature into a resource for experimental analysis and discovery

**DOI:** 10.64898/2026.05.04.722753

**Authors:** Lukas Oesinghaus, Molly Park, Rulin Shao, Pang Wei Koh, Georg Seelig

## Abstract

Cytokine biology is dispersed across hundreds of thousands of publications, making it difficult to use systematically when interpreting new experiments. Large language models (LLMs) can assist with focused literature interpretation, but ad hoc retrieval remains incomplete and unreliable. We present the Cytokine Effect Database (CytED), a framework for interfacing user-supplied experimental datasets with literature knowledge at scale. CytED uses a multi-step LLM pipeline to generate over a million cytokine-cell type-effect triples from 110,000 full-text publications, with annotations for experimental context and directional changes in genes, pathways, and cellular processes. This structure enables quantitative comparison between observed perturbation responses and prior literature across cytokines, cell types, and experimental contexts. Applied to in vitro IL-10 stimulation of PBMCs, CytED identifies unexpected pro-inflammatory features in monocytes and systematic in vivo-in vitro differences in cytotoxicity responses in CD8+ T cells. CytED infers cytokine signaling, distinguishes primary from secondary cytokine effects, and guides the design of combinatorial perturbation screens. Together, CytED establishes a general paradigm for converting unstructured domain literature into analytical tools that bridge literature and experiment.

## Introduction

Cytokines are central regulators of immune cell state and function, acting in highly context-dependent and pleiotropic ways^1^. Decades of research have generated an enormous and rapidly expanding body of literature: PubMed lists over one million cytokine-related publications, and even a single cytokine such as IL-12 is associated with more than 30,000 papers (Methods). Advances in high-throughput technologies now enable systematic perturbation and multimodal measurement of cytokine responses at unprecedented scale, promising to accelerate progress in disentangling their pleiotropic effects^2–5^. This promise depends on effective integration with prior knowledge, because it is otherwise challenging to properly contextualize the observed response patterns. However, relevant information is buried across tens of thousands of heterogeneous publications, which necessitates tools that enable scalable interaction with this existing knowledge.

Prior efforts to systematize the cytokine literature have taken several forms. Curated databases, whether manually assembled^6,7^ or constructed using classical natural language (NLP) processing approaches^8,9^, provide structured summaries but face inherent trade-offs: manual curation cannot keep pace with the volume of publications, while classical NLP methods struggle to capture nuances in natural language. Neither approach readily interfaces with high-dimensional experimental data. Alternatively, transcriptomic datasets^10,11^ enable systematic comparisons or summaries across cytokine treatments. While powerful for some applications, this strategy misses the causal inferences provided by publication text.

Large Language Models (LLMs) provide a novel approach to interfacing scientific literature. They can retrieve scientific knowledge^12^, produce encyclopedia-like summaries^13^, propose novel scientific hypotheses^14,15^, assist in experimental data evaluation^16^, and even perform end-to-end scientific workflows^17^. However, these approaches are constrained by practical limits on the considered corpus size and challenges in ensuring reproducibility, factual reliability, and human legibility^18^. Alternatively, LLMs can parse the literature at scale to generate scientific knowledge bases, which has previously been applied in materials science^19,20^, pathogenic variant annotation^21^, and immune cell type-specific knowledge graph construction^22^. However, most such efforts have been limited to the database construction step without modeling the causal structure or directly interfacing with novel experimental data.

Here, we present the Cytokine Effect Database (CytED), a repository of 1,012,726 statements of the form ‘cytokine x has effect y on cell type z’ extracted from 110,000 publications using multi-step processing by an LLM reasoning model (DeepSeek-R1, **Fig. 1a**)^23^. Each such statement is annotated with affected genes, pathways, and cell processes as well as metadata on used experimental conditions, such as cytokine concentrations or timing of the experiment. Extraction quality is ensured by a four-factor quality control strategy (QC) consisting of two QC metrics generated during extraction and processing and two dedicated QC filtering steps. After describing the construction of CytED, we first demonstrate its use in analyzing the cytokine literature itself. We generate “textbook” consensus statements, quantify and investigate the experimental context dependence of cytokine effects, and compare cytokine similarities across cell types. We then apply CytED to the analysis of experimental datasets. We use CytED both to quantify general literature-experiment agreement and to support legible interpretation of cytokine perturbation responses in key signature genes. For example, CytED shows that IL-10 stimulation of PBMCs produces the expected myeloid deactivation program, but also highlights context-specific pro-inflammatory deviations in monocytes and missing cytotoxicity responses in CD8 T cells. By retrieving a small, interaction-specific set of literature passages together with their metadata, CytED constrains LLM-based synthesis to evidence that is highly relevant and limited enough for direct inspection. We then generate literature-based gene sets that rival a gold-standard approach trained on aggregated experimental data^11^ in identifying cytokine signaling and find associations between cytokine signaling and cellular processes in tumor datasets^24,25^. We derive literature priors to distinguish primary and secondary effects in perturbation datasets. Finally, we propose a literature-informed high-throughput screen of combinatorial cytokine perturbations.

**Fig. 1.**
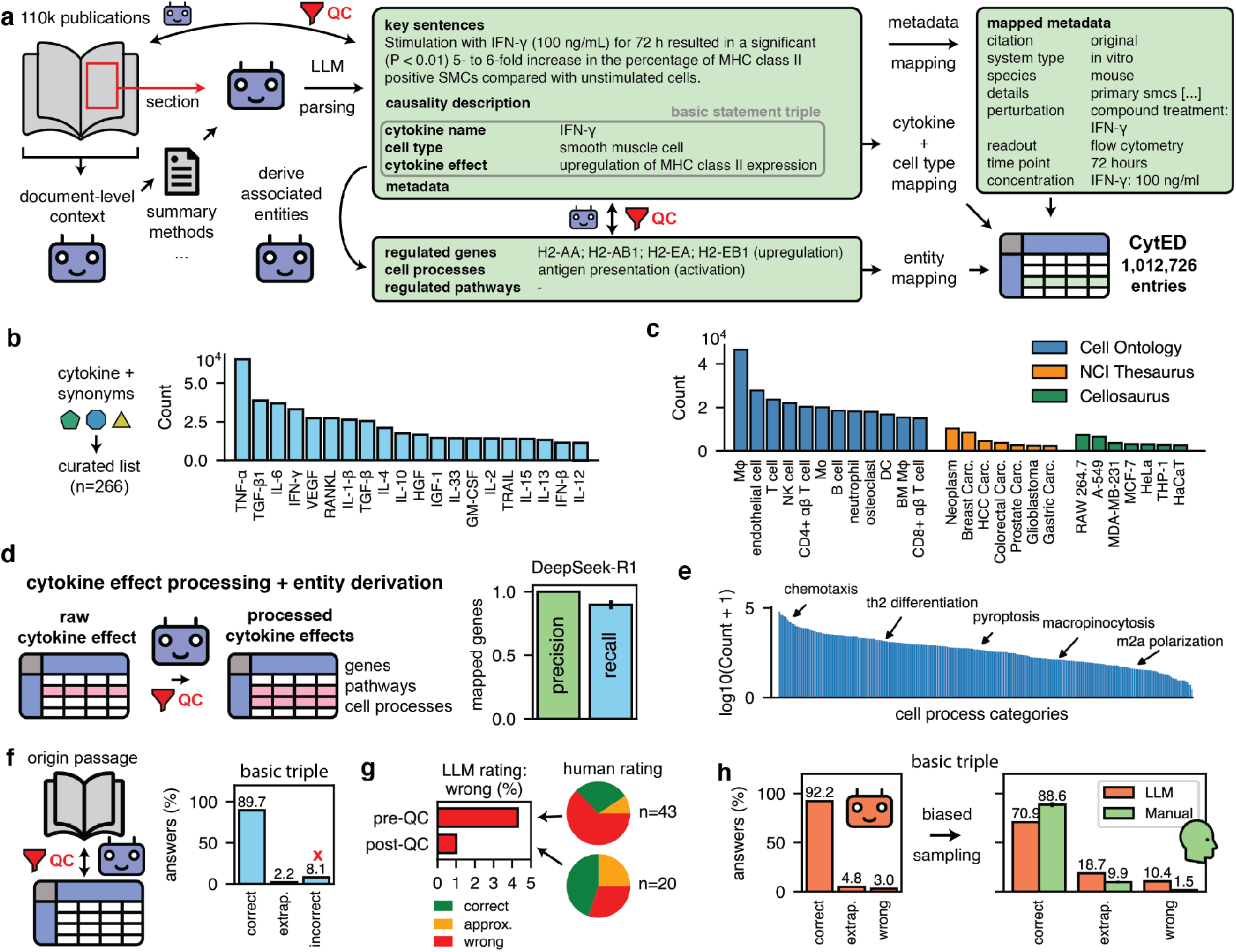
Processing the cytokine literature into a database using large language models. **a**, Overview of the construction workflow for the Cytokine Effect Database (CytED) for a single example row. **b**, Cytokine names were standardized using a manually curated list of target cytokines. The bar graph shows the cytokine counts from 110,000 papers for the most common cytokines. **c**, The most commonly mapped cell types for each target ontology. **d**, Initial cytokine effect entries are processed into one or, if the initial entry mixes multiple concepts, multiple output entries containing genes, pathways, and cell processes. The bar graph on the right shows precision and recall of DeepSeek-R1 for gene-level statements for a test set of 52 manually processed input rows (117 output rows). **e**, Distribution of literature counts for mapped cell process categories. **f**, The initial parsed data undergoes quality control by a DeepSeek-R1-powered comparison between the original text chunk and the extracted table entries. Incorrect entries are removed from the table. **g**. Table entries pre- and post-QC were evaluated for the presence of clear errors by GPT-5.2 (n=1000 and n=2000, respectively). Incorrect entries were then blindly re-rated by a human evaluator. **h**, The filtered table was re-evaluated according to the same QC prompt as in (f) by GPT-5. The resulting entries were sampled to increase the expected error rate and presented to human evaluators.

CytED serves two primary purposes. First, it is a reference and analytical resource for immunologists. We provide a web interface (https://immune-context.github.io/cyted/) to allow researchers to quickly find relevant information and a software package (pyCytED) to support the analysis of cytokine signaling in experimental datasets. Second, it is an end-to-end framework that demonstrates how LLM-extracted knowledge bases can enable literature-grounded high-throughput interpretation of experimental datasets and follow-up experimental design. Thus, while CytED itself is constructed to capture cytokine effects, it also represents a more general, scalable paradigm for using LLMs to connect high-throughput experiments and an ever expanding literature. We provide a software package for literature database construction (papers2db) to encourage researchers to generate similar tables for their particular purpose.

### LLM-parsing of the cytokine literature creates a repository of non-standardized statements

We used EuropePMC^26^, which contains around 6 million articles, as our open-access publication repository. To find relevant publications, we first searched PubMed using manually curated cytokines (e.g., IL-17A), cytokine isoforms (e.g., IL-32β), and cytokine families (e.g., Type I IFNs) for a total 266 unique cytokine entries and their curated synonyms (**Fig. S1a, Table S1** and **S2**). We then combined LLM-derived (Gemini-2.0-Flash) relevance ratings (**Fig. S1b**), age-adjusted citation count percentiles (**Fig. S1c**)^27^, and a cytokine diversity metric to rank publications (**Fig. S1d-e**, Methods). The algorithm was applied separately for original publications and reviews. We picked the top 100,000 original publications and top 10,000 reviews (**Fig. S1f**).

The initial LLM-based parsing extracts statement triples of the form ‘cytokine x has effect y on cell type z’ and aggregates them into a table (**Fig. 1a, Fig. S2**, see **Table S3** for all prompts used in this publication). Thus, each row consists of one or more cytokine names, cell types, and cytokine effects. For each row, we first extracted key sentences to keep statements traceable to specific lines in a publication and a description of the overall causality, e.g., ‘IFNγ upregulated MHCII in smooth muscle cells’. We then extracted the desired information and augmented it with rich metadata, amongst others: the experimental system (species; in vitro or in vivo experiment; other details); applied experimental perturbations (e.g., genetic knockouts); used readouts (e.g., flow cytometry); time points (e.g., 24h); concentrations; whether the statement is original to the publication or cited from another and if so, the identity of the origin publication. Finally, the LLM was also prompted to generate an integer confidence score between 1 and 10 measuring its level of trust in its own extraction.

This initial task is highly complex. The LLM must follow the formatting requirements for each entry column and correctly associate each statement with its experimental metadata. Most challengingly, it must understand the causality of a given statement, for example, that upregulation of gene A in a receptor knockout for a cytokine B implies ‘cytokine B downregulates gene A’. Smaller models, e.g., DeepSeek-R1-Distill-Qwen-32B, often failed catastrophically at this task and even proprietary non-reasoning models (e.g., Gemini-2.5-Flash, GPT-4.1) still frequently struggled (**Fig. S3a**). On the other hand, the best reasoning models available at the time of CytED construction (e.g., GPT-5) performed well but were too expensive for our desired scale. We found that DeepSeek-R1 (∼30 times cheaper than GPT-5), a near-state-of-the-art reasoning model at the time, had an excellent trade-off between cost and performance (**Fig. S3a**). Each publication was chunked into sections according to publication subheadings to strike a balance between parsing cost, sufficient context, and a decay in extracted information for longer inputs (**Fig. S3b**). Because individual text sections can lack necessary metadata, we decorate it with summarized document-level information (**Fig. 1a, Fig. S2**). Across the 110k parsed publications, DeepSeek-R1 extracted a median of 9.2 statements per publication. Notably, parsing abstracts yielded only 0.67 statements per publication.

### Entity mapping and post-processing generates a unified database of cytokine effects

To standardize cytokines, we used our curated list of names and synonyms to generate name variants mapping to the 266 standard cytokine entities. The most common mapped cytokines are TNFα, TGFβ, and IL-6, although almost all cytokines are present to some degree (**Fig. 1b, Fig. S4**). Cell types are mapped to established ontologies (Cell Ontology^28^, NCI Thesaurus^29^, Cellosaurus^30^) using a combination of LLM standardization and classification, similarity-based candidate retrieval, and LLM choice (Methods). We verified the mapping algorithm using 440 manually mapped entries, where we observe about 1% off-target mapping, e.g., ‘metaplastic epithelial cell’ to ‘epithelial cell’ instead of ‘metaplastic carcinoma’ (**Fig. S5**). The most common cell ontology entries are macrophage, endothelial cells, and T cells, the most common NCI Thesaurus entries are neoplasm, breast carcinoma, and hepatocellular carcinoma, and the most common cell lines are RAW 264.7, A-549, and MDA-MB-231 (**Fig. 1c**). Overall, however, mapped cell types are highly diverse (3653 unique), and the top 50 only represents around 57% of all statements.

Cytokine effect entries, i.e., free-text statements extracted by the LLM about the impact of a cytokine in a particular context, were processed into one or more logically distinct sub-effects using LLM calls, e.g., ‘activation of chemotaxis and cytotoxicity’ into ‘activation of chemotaxis’ and ‘activation of cytotoxicity’. Each processed statement is associated with zero or more affected genes, pathways, and cell processes, i.e., cell-level behavior such as chemotaxis, by the LLM, each with a defined direction of change (**Fig. 1d**), then standardized, enabling computational analysis based on these standardized entities. We also added a ‘causality judgement’ that asks the LLM to evaluate whether the provided key sentences and causality description are internally consistent with the mentioned cytokines, cell types, and cytokine effects. To map genes, we rely on the LLM’s ability to associate free-text statements with genes that are either explicitly mentioned or strongly implied. For example, ‘MHC class II expression’ in mice plausibly implies upregulation of *H2-Aa, H2-Ab1, H2-Ea*, and *H2-Eb1*. For manually curated test data, DeepSeek-R1 again presents a good balance between performance and cost with almost 100% precision and 90% recall and was therefore chosen as the working model (**Fig. 1d, Fig. S6**). Cell processes are mapped to a manually curated list of 263 process types using an approach similar to what was employed for cell type mapping (**Fig. 1e, Table S4**), yielding a range of common to rare statements, e.g., chemotaxis (16,501 entries), pyroptosis (471 entries), or m2a polarization (41 entries). Pathways were similarly mapped to the pathway ontology^31^ (**Fig. S6e**).

Metadata columns were standardized using a variety of approaches customized to the particular properties and distribution of entries in each column (**Fig. S7**, Methods), yielding for example, a 50/50 mix of cited and original statements in original publications (**Fig. S7a**), an approximately equal number of statements for humans and mice, and four times as many *in vitro* as *in vivo* statements (**Fig. S7c**). Immunoblotting, qPCR, and flow cytometry are the most common experimental readouts (**Fig. S7d**), SARS-CoV-2 is the most common viral challenge (**Fig. S7e**), and 24h treatment with 10 ng/ml is the most common condition for exogenously supplied TNFα (**Fig. S7f-g**).

### A four factor quality control strategy reduces the error rate

Although the initial extraction quality was generally good, it was still easy to manually find errors such as inverted causality. To reduce the error rate, we performed two dedicated rounds of QC to supplement the confidence score of the initial extraction and the causality judgement of the reformulation step, giving the entire QC process four total stages. In the first round, each row is presented to DeepSeek-R1 along with its original publication text for evaluation. The prompt was structured to find any entries containing even mildly incorrect information in its primary entries (cytokine, cell type, cytokine effect), resulting in a relatively high fraction of 8.1% of entries rated as incorrect (**Fig. 1f**). As we valued precision over recall, we simply removed these entries from the table. For experimental metadata, entries that were labeled as incorrect were removed without changing the rest of the row. In the second dedicated QC round, we applied a similar approach to reduce the error rate of the effect reformulation step (cf. **Fig. 1d**). The reformulation step has a lower error rate but QC still alters at least one column in 2.4% of rows and correctly removes, e.g., manually detected incorrect double negatives in the cell process direction (**Fig. S8a-b**).

Taken together, the QC process strongly reduces the overall error rate. Using GPT-5.2 as a judge, we find a reduction from 4.3% to 1.0% in entries marked as having clear errors in the basic statement triple from pre-to post-QC (**Fig. 1g**). Blinded manual inspection of these LLM-labeled errors finds that while 63% of pre-QC LLM-labeled errors are genuinely incorrect, often due to incorrect causal relationships, only 30% of post-QC LLM-labeled errors are genuinely incorrect, and these entries are largely free of obvious causal errors. Thus, the rate of clear errors is reduced approximately 9-fold by the QC process.

To estimate an upper bound for the error rate of the final quality-controlled database, we first re-evaluated 321 publication sections (1348 rows) of the table using GPT-5, finding a 3% LLM-labeled error rate (**Fig. 1h**). We then subset to publication chunks containing at least one statement labeled as either incorrect or approximate, thereby biasing the manual evaluation towards finding an upper limit for the true error rate. Two human evaluators rating 134 such entries rate a mean of 1.5% of entries as containing genuine errors (**Fig. 1h**). For metadata columns, we find a mean error rate of 0.7% (**Fig. S8c**).

The entity-mapped and quality-controlled table of cytokine effects with metadata constitutes the Cytokine Effect Database (CytED), containing 1,012,726 annotated effects for individual cytokine-cell type pairs.

### CytED maps the literature landscape of cytokine regulation

In total, CytED contains 167,299 unique cytokine-cell type-gene interactions covering 7,670 unique genes, 164,656 unique cytokine-cell type-cell effect interactions, and 41,424 unique cytokine-cell type-pathway interactions (**Fig. S9a-c**). The most common gene-level statements all reflect well-established canonical facts, such as IL-12-induced production of *IFNG* in natural killer cells (NK cells)^32^, RANKL-induced upregulation of *NFATC1* in bone marrow macrophages (BM MΦ)^33^, or IL-10-induced downregulation of TNF in macrophages^34^ (**Fig. 2a**). Common cell process or pathway statements similarly converge on well known facts, e.g., angiogenesis in endothelial cells due to VEGF^35^ (**Fig. 2b**), or inhibition of RANK signaling in osteoclasts by osteoprotegerin^36^ (**Fig. S9d**). Using the example of dendritic cells (DC), we can quickly retrieve a ranked list of cytokines that are discussed in the literature as upregulating DC cell chemotaxis (**Fig. 2c**) with canonical lymph node homing via CCL19 and CCL21 being most prominent^37^.

**Fig. 2.**
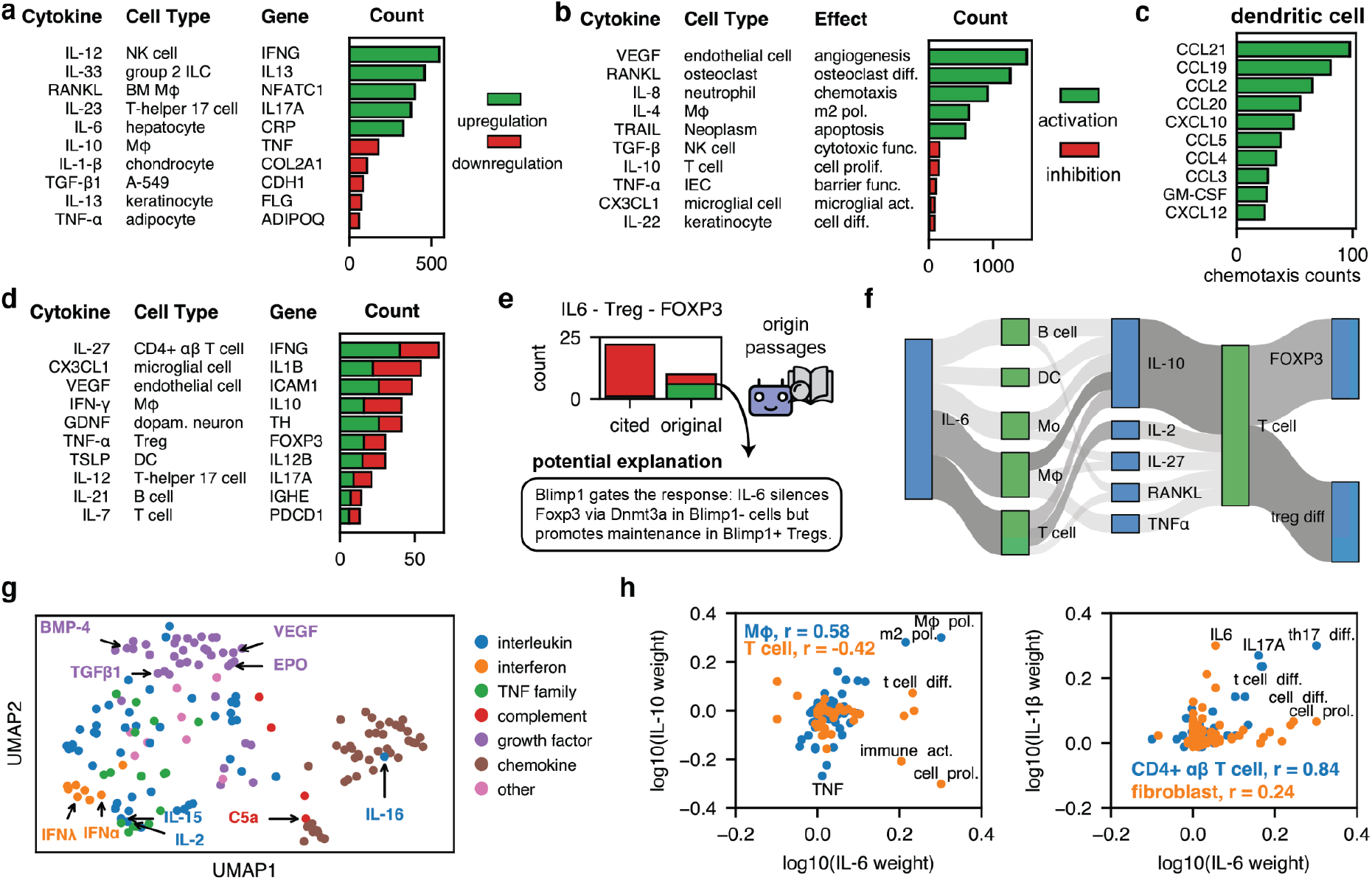
Systematic investigation of the cytokine literature using CytED. **a**, Publication count for some of the most common cytokine-cell type-gene regulation statements. **b**, Publication counts for the most common cytokine-cell type-cell effect statements. **c**, Top cytokines that activate chemotaxis in dendritic cells. **d**, Up- and downregulation counts for the most common cytokine-cell type-gene regulation statements with a low directional agreement. **e**, The link between CytED and literature fragments was used to formulate potential explanations of a divergent distribution of directions for the effect of IL-6 on FOXP3 expression in Tregs between cited and original statements using LLM queries. **f**, Downstream cytokines upregulated by IL-6 that upregulate FOXP3 or activate the Treg differentiation process. The shading of each connection indicates log2+1 of relative literature counts and thereby the prominence of a connection. **g**. UMAP of different cytokines for log10+1-transformed net literature counts of merged cell type-cell effect and cell type-gene relationships after truncated singular value decomposition. Functionally similar cytokines group together. **h**. Literature weights of different cell processes and genes for the effect of (left) IL-6 and IL-10 in macrophages (MΦ) or T cells and (right) IL-6 and IL-10 in fibroblasts and CD4+ αβ T cells. The Pearson correlation r is shown as an inset.

Most genes and cell processes are near exclusively either up- or downregulated in a given cell type in response to a given cytokine, but about 6% of all statements with at least 6 reported publications show below 80% concordance in their direction (**Fig. S10**). For the effect of TNFα on *FOXP3* expression in T regulatory cells (Treg)^38^ or IL-27 on *IFNG* expression in CD4 T cells^39^ up- and downregulation statements are nearly equally common (**Fig. 2d**). This lack of concordance does not imply direct disagreement between different literature reports but the context-sensitivity of cytokine effects captured by CytED.

In its metadata, CytED distinguishes between original research findings and claims based on cited literature. Interestingly, the consensus of cited statements sometimes seemingly disagrees with reported original research findings: For the effect of IL-6 on the key differentiation marker *FOXP3* in Tregs, we find near universal downregulation in the secondary literature, making IL-6 a textbook antagonist to Treg differentiation and maintenance^40^ (**Fig. 2e**). In the original research papers within our corpus, the claimed effect direction is much more heterogeneous. We used CytED to retrieve relevant text passages and presented it to GPT-5.2 to generate potential explanations of this discrepancy. One of the retrieved research papers is highlighted as discussing how the induction of Blimp1 in inflamed tissues prevents downregulation of *FOXP3* under IL-6 stimulation^41^, and that this is the experimental context of the publications that find upregulation of FOXP3 by IL-6. This implies experimental context as a partial explanation, namely for a lack of downregulation, though by itself not actual upregulation.

Because regulated genes often code for cytokines, CytED can be converted to a directed, weighted graph in which cytokines, genes, and cell effects are connected by up- and downregulation statements via cell types, with edge weights being provided by literature counts. This, in turn, allows higher-order effects to be quantified. Returning to the question of why IL-6 might upregulate FOXP3 in some cases, we checked whether IL-6 can release secondary cytokines capable of driving *FOXP3*/Treg differentiation in T cells. Upregulation of IL-10 in macrophages and T cells and of IL-2 T cells emerge as the most well-established connections (**Fig. 2f**), yielding two testable hypotheses for the inversion of the effect of IL-6 in some experimental datasets.

We used literature counts to create a vector of cell type-gene and cell type-process counts for each cytokine, counting downregulation counts as negative (Methods). Signed log2+1 conversion then yields a signature vector for each cytokine, which we used to investigate the global landscape of cytokine function. We generated a two-dimensional embedding of these signatures to visualize functional cytokine groupings (**Fig. 2g, Fig. S11**). In this landscape, canonical and de facto chemokines (e.g., C5a and IL-16) form a separate cluster, validating functional embedding. Other cytokines also show expected patterns: IFNα is close to IFNλ, IL-2 to IL-15, and BMP-4 to TGFβ1. Some cytokines have highly divergent functions in different cell types, information that is obscured by this more global view but can easily be extracted from CytED: Literature counts per cell effect for IL-6 and IL-10 are positively correlated (r=0.58) in macrophages due to their shared tendency to cause M2 polarization, while in T cells, they are negatively correlated because IL-6 causes immune activation while IL-10 dampens immune responses (**Fig. 2h**). Similarly, IL-6 is similar to IL-1β in CD4+ αβ T cells, where they act as co-factors for Th17 differentiation (r=0.84), while in fibroblasts, their functions barely correlate (r=0.24), with IL-6 causing a proliferative response and IL-1β causing an inflammatory response.

In summary, CytED provides a framework for quantifying canonical cytokine interactions while also systematically surfacing the biological discrepancies that exist across the literature. Further, because entries are anchored to original research excerpts, CytED enables efficient and highly targeted LLM-powered literature mining on specific questions. Finally, the overall pattern of annotated relationships quantifies systematic relationships between cytokines across cell types.

### CytED is a structured interface for the analysis of experimental results

The primary purpose of CytED is to serve as a tool to interface new experimental datasets with the existing literature, providing useful priors for experimental analysis (**Fig. 3a**). In the following, we will demonstrate how CytED allows for data exploration based on shared cytokine-cell type-gene effect triples. We started with a recent scRNA-seq dataset of 90 cytokine perturbations of human PBMCs containing 12 major cell types (**Fig. 3b**). We matched CytED cell type entries to experimental cell types using the cell ontology (**Fig. S12a-c**), leading to 7,764 unique shared cytokine-cell type-gene regulation triples between CytED and the experiment, where each triple can be evaluated for its directional agreement between literature observations and experimental data. We coarse-grained experimentally observed directions of change in gene expression using the observed log2 fold change (log2FCs) with threshold of 0.5, resulting in around 2% of all triples being classified as up- or downregulation each (**Fig. S12d**). We find good agreement in triples that are observed in both experiments and literature (**Fig. 3b**), with 45% shared direction, 6% opposite directions and 49% without significant experimental changes. We note that it is not surprising that many triples are observed in the literature, which captures a wide range of contexts, but not in a particular screen with highly specific experimental conditions. Within-cell type shuffling of experimental cytokine labels reduces this to 16% shared directions, 7% opposite directions, and 77% without significant regulation in the experiment. Higher shared than opposite directions in the shuffled case likely reflect similar regulation for many of the strongest cytokines in the screen (e.g., IFNβ and IFNγ).

**Fig. 3.**
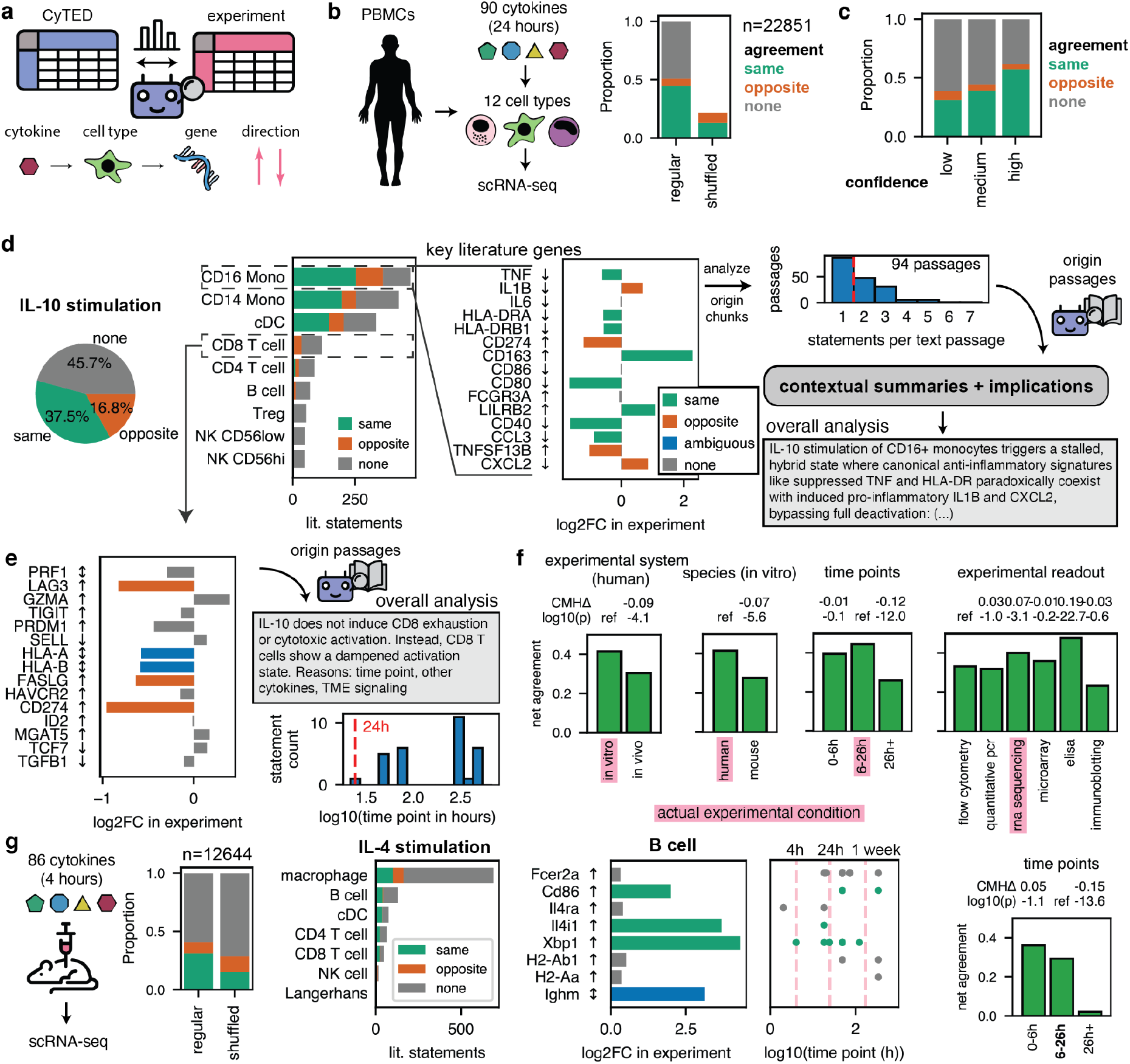
CytED identifies signatures of cytokine signaling in experimental datasets. **a**, CytED entries and transcriptional data are merged by matching cytokine-cell type-gene triplets. **b**, A high-throughput in vitro screen of human PBMCs response to treatment with 90 cytokines is used as a primary test case^4^. The left bar graph shows the agreement between CytED statements and the experimental data. The right bar graph shows agreement after shuffling cytokine labels within each cell type as a control. **c**, Literature-experiment agreement by triple confidence derived from literature abundance. **d**, Workflow to analyze experimental changes in key genes, illustrated for IL-10 stimulation of CD16+ monocytes. For each cell type, a subset of experimental changes agrees or disagrees with CytED entries. Key literature genes for IL-10 stimulation of CD16+ monocytes are shown in the inset. Each triple can be traced back to specific places in origin publications. The experimental results and corresponding text passages are provided to an LLM to generate contextual summaries. Summaries across all key genes are then combined to produce an overall interpretation of the experimental response. **e**, Same analysis as in (d) for IL-10 stimulation of CD8+ T cells. The distribution of experimental time points for statements is shown on the bottom right. **f**, Dependence of the agreement (same-opposite) between experimental data and CytED on metadata variables: experimental system (human studies only), species (in vitro studies only), measurement time point, or experimental readout. Statistical significance was determined via a generalized Cochran-Mantel-Haenszel (CMH) test stratified by cytokine identity relative to the reference group (ref.), with the weighted mean score difference (CMHΔ) as the effect size. Multiple testing was corrected using the Benjamini-Hochberg procedure. **g**, Analysis as in (d) and (f) for a 4-hour in vivo cytokine stimulation experiment in mice. General agreement is much lower, at least partially due to higher noise levels in the data. Overall, we find that literature entries from long term experiments are overwhelmingly less likely to agree.

We reasoned that more common statements in the literature should be more likely to generalize, reflecting their function as key marker genes for a given cytokine perturbation. We assigned low, medium, and high confidence labels based on how often a statement occurs in the literature (**Fig. S13a**) and tested agreement with the PBMC perturbation dataset. Higher confidence statements are much more likely to agree with the experiment (**Fig. 3c, Fig. S13b**). For IL-10 treatment, 41% of statements in CytED have the same direction as what is observed experimentally, 41% show no change, and 18% display a change in the opposite direction. On a per cell type basis, agreement is relatively high for CD16 monocytes, whereas for CD8 T cells, the experiment overwhelmingly shows no or opposite responses. These agreement fractions are dominated by the aforementioned key genes, which in CD16 Mono, core literature responses include downregulation of *TNF* (observed), *IL1B* (opposite is observed) and *IL6* (no change) (**Fig. 3d**).

CytED links the observed changes in key genes to particular publication passages, thereby functioning as a structured interface for automated LLM-based analysis (**Fig. 3d**). This makes it possible to scale literature-grounded interpretation across many cytokine-cell type conditions, without sacrificing human interpretability or clearly traceable source attribution. We presented GPT-5.2 with all text passages that discussed two or more key genes for the effect of IL-10 on CD16 monocytes (94 text passages), and asked it to summarize and analyze this chunk in light of the particular changes observed in the provided experimental data. The results of this per passage summarization and analysis were then used together with all experimental results for key genes to yield an overall analysis. The model-generated summary suggested ‘an intermediate or heterogeneous cell state that combines features of IL-10-driven reprogramming with retained or emergent pro-inflammatory potential’, showing how CytED helps prioritize a small set of key genes, recover the most important literature-supported aspects of a response program, and expose specific divergences, such as TNF and IL1B, for a more detailed follow-up. In CD8 T cells, core literature responses to IL-10 include upregulation of *LAG3* (opposite is observed), *HAVCR2* (no change), and *GZMA* (no change), genes associated with exhaustion and cytotoxicity (**Fig. 3e**). LLM-based literature passage analysis therefore notes that in the experiment ‘IL-10 does not induce CD8+ T cell exhaustion or cytotoxic activation’, noting the time point, in vivo context, or tumor microenvironment signaling in the source publications as potential reasons. The lesson here is different: CytED indicates that this perturbation context may not be well suited to studying direct in vivo-like IL-10 signaling in CD8 T cells, and is instead consistent with the possibility that observed CD8 T cell effects could be secondary and mediated by myeloid cells.

Many of these potential reasons for disagreement between the literature and the experiment are present as metadata in CytED. For example, time point metadata for IL-10 treatment of CD8 T cells indicates that nearly all studies were conducted using time frames of at least 48h and usually well over a week, which by itself plausibly resolves literature-experiment disagreement in this case (**Fig. 3e**). Using a meta-analysis stratified by cytokine identity, we asked whether metadata similarity to the studied experiment globally influences the likelihood of agreement with the experimental data (**Fig. 3f**). Entries from in vitro studies are more likely to agree than those from in vivo studies (padj=10^-4.1^), and so are entries from human studies (padj<10^-5.6^), and short term studies (<26h) (padj<10^-12^). The measurement modality also plays a role, with triples from RNAseq more likely to agree than those derived from flow cytometry (padj<10^-3.1^). Thus, a closer match in experimental conditions translates to higher agreement for this study. A similar analysis lends support to the QC steps (**Fig. S14**). We found a linear increase in the net agreement with the initial parsing confidence score, going from 0.25 to 0.4 for a confidence score of 5 and 10. Similarly, agreement is higher for statements that passed the causality check of the reformulation step (padj=10^-44.9^), and for statements that passed the first QC filtering step (padj=10^-34.3^).

The same workflow presented here is straightforwardly applicable to other experimental datasets. In an *in vivo* screen where mice were stimulated with cytokines for 4 hours^2^, overall agreement with the literature is lower at 31% shared and 10% opposite directions versus 17% the same and 14% the opposite direction for shuffled labels (**Fig. 3g**), presumably partially due to differences in the experimental setup and partially due to much higher noise levels in the associated scRNA-seq pseudobulks resulting from smaller cell numbers (**Fig. S12e**). For IL-4 stimulation, the B cell response in this experiment contains some key genes that respond as expected, such as *Cd86, Xbp1, Ighm*, and *Il4i1*, while many other canonical genes do not. Looking at metadata, we find that, in this case, the time point does not cleanly explain the differences. However, across the entire experiment, we do find a striking pattern of long term studies (>26h) being overwhelmingly less likely to agree (padj=10^-13.6^), consistent with the short time frame of the experiment.

### CytED-derived gene sets correctly infer cytokine signaling activity

The signature of a cytokine perturbation of a given cell type is characterized by a quantitative pattern of gene up- and downregulation. Knowledge of such signatures enables identification of candidate cytokines that drive such differences^11^. The question of which cytokine likely produced an observed response is relevant in the common situation where gene expression changes are induced via cytokine signaling triggered by a biological event such as a viral infection that is not simply the external addition of a known cytokine. We wondered whether we could use CytED to generate such quantitative cytokine signatures and thus perform such inference. To generate quantitative signatures, we aggregated gene counts across cell types for all cytokines, counting upregulation as positive and downregulation as negative counts, and used a signed log2+1 conversion to generate weighted gene sets. Signaling activity is inferred using a univariate linear model^42^ on log2FCs (**Fig. 4a**).

**Fig. 4.**
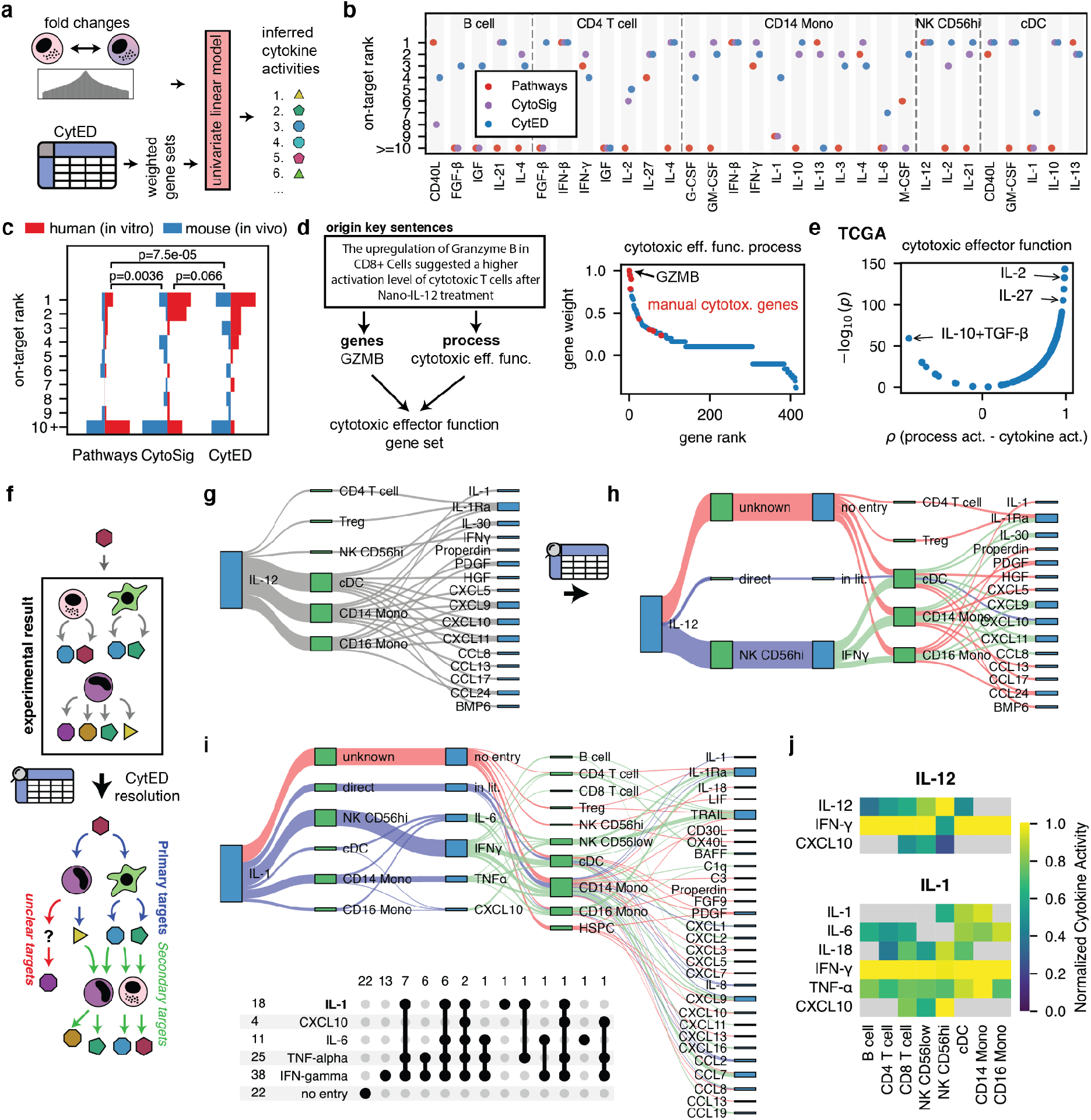
CytED is a platform for the analysis of cytokine signaling. **a**, Count distributions of cytokine-cell type-gene pairs are converted to weighted gene sets to infer cytokine signaling activities via a univariate linear model. **b**, Ranking of the on-target cytokine for pathway-derived, CytoSig, and CytED-derived weighted gene sets in a subset of perturbation conditions for human PBMC data. **c**, Distribution of ranks as in (b) for both human PBMC and in vivo mouse cytokine perturbation data. The p-value is calculated using a two-sided Wilcoxon signed-rank test on paired on-target ranks across datasets. **d**, Gene sets for specific cell processes are generated by identifying genes that co-occur in the same origin row. Co-occurring gene counts across CytED then yield a gene signature for a given process, here shown for cytotoxic effector signaling. Manually curated cytotoxicity genes^25^ (CD8A, GZMA, GZMB, GZMH, GZMK, PRF1, NKG7) are highlighted in red. **e**, Spearman correlation between inferred cytotoxic effector function process activity and the activities of different cytokines in melanoma samples from the TCGM dataset. **f**, Perturbation experiments measure a mixture of primary and secondary effects. Using CytED, one can infer which effects are likely primary, which secondary, and which might be novel cytokine effects. **g**, Experimental results of IL-12-cell-cytokine regulation connections in the human PBMC dataset. Each connection indicates a connection with log2FC>1 and padj<0.05. **h, i**, Inference of primary and secondary effects of (h) IL-12 and (i) IL-1 based on CytED literature annotations. The inset shows how many connections are explained by the primary cytokine, secondary cytokines, or are not annotated in CytED for any of the cytokine perturbations. **j**, CytED-based cytokine activity analysis for IL-12 and IL-1 stimulation.

Various other approaches have previously been used to identify cytokine signaling. One coarse-grained approach is to look for transcription factor activities in core cytokine pathways, e.g., STAT6 for IL-4^7,43^. A more powerful approach is to train predictive models on perturbation datasets^11^. We therefore use manually curated core pathway transcription factors (**Table S5**) in combination with CollecTRI^44^ for transcription factor activity as a relatively easy-to-beat benchmark and the predictive model CytoSig^11^ as a hard-to-beat benchmark. We first tested performance on perturbed primary cells in the CytoSig dataset/training data itself (n=507, **Fig. S15a**), placing the correct cytokine in the top 3 in 66% of all cases, compared to 51% for pathway-derived gene sets and 85% for CytoSig itself. Performance varies across cytokines, with, e.g., TNFα gene sets performing well and TGFβ gene sets much more poorly, possibly because the typical experimental context of TGFβ in the immunological literature is more distant from in vitro perturbation studies (**Fig. S15b**).

As a first independent test set, we chose cytokines with strong observed, likely primary signaling activity in the human PBMC experimental dataset^4^ (**Fig. 4b**, Methods, **Table S6**). Both CytoSig and CytED gene sets consistently outperform pathway genes, placing the correct cytokine on rank 1 to 3 in 21 and 22 out of 31 cases, respectively. CytED outperforms CytoSig more often than not on this dataset (12 wins, 11 draws, 8 losses). To verify that this is not particular to this dataset, we used a second test set from the aforementioned scRNA-seq dataset of in vivo cytokine perturbation treatments of mice (**Fig. S15c, Table S6**). For this dataset, the hit rate for cytokine activity inference is in general worse, probably again due to the more complicated in vivo environment and noisier pseudobulks. Still, we again find that CytED outperforms CytoSig (12 wins, 8 draws, 6 losses). Thus, literature-derived CytED cytokine signatures rival or outperform perturbation-trained signatures on both of these datasets (overall p=0.066, **Fig. 4c**). CytED also contains gene sets for many cytokines that are currently missing in CytoSig (∼124 in CytED for non-combinatorial signatures with at least 20 different genes versus 51 in CytoSig). However, there are many individual cases where CytoSig outperforms CytED gene sets, and some gene sets perform better than others. As their information derives from largely non-overlapping sources, namely high-throughput transcriptional perturbation data for CytoSig and curated genes from individual experiments for CytED, it would likely be optimal to combine their gene sets for a consensus analysis, where both are available.

### CytED identifies core processes and cytokine signaling in tumor data

Recent work identified IL-27 as promoting cytotoxic CD8+ T cell responses in tumors. This effect was initially found by correlating the presence of cytotoxicity genes with *IL27* expression in melanoma samples from The Cancer Genome Atlas (TCGA)^25^. Reproducing this initial analysis, we find a Spearman correlation of 0.74 (**Fig. S16a**), though *IL27* is far from having the strongest association.

We first asked whether we could use CytED to generate gene sets for specific processes that can be used to look for process activity analogous to the inferred cytokine activity in the previous section. To do so, we count processes and genes that derive from the same origin row as associated and count co-occurence (**Fig. 4d**). For example, if the same statement leads to the annotation of *GZMB* upregulation and also cytotoxic effector function process activation, it is reasonable to assume that the two are associated. For the cytotoxic effector function process, this yields a gene set that prominently includes all of the manually curated cytotoxicity genes, amongst many others. Looking again at the correlation with IL-27 expression, we find near perfect correlation with the results from using the mean of manually curated cytotoxicity genes (**Fig. S16b**, ρ=0.99). Next, we associated cytotoxicity process activity with cytokine signaling activity. For the inferred IL-27 activity, we see ρ=0.97, a much cleaner signal than for comparison with cytokine gene expression. This is not definitional as the correlation between the weighted gene sets for IL-27 activity and cytotoxicity is only 0.23 (**Fig. S16c**). Other top hits (IL-2, IL-15) are also likely causal for cytotoxic T cell activity, indicating that using cytokine signaling instead of gene expression yields a much better prior for causal associations than gene expression.

To further validate our process gene sets, we looked at inferred process activities across tumor cell types in TCGA (**Fig. S16d**). We find that angiogenesis is most strongly enriched in kidney clear cell carcinoma^45^, net formation in lung adenocarcinoma^46^, and immune response activity shows the lowest enrichment in uveal melanoma^47^, all of which agrees with known biology. We performed a within-cancer-type metanalysis of the association between cytokine activities and process activities after regressing out leukocyte fractions within each tumor type. We found that TGFβ is strongly associated with fibrosis activity, VEGFA with angiogenesis activity, and IL-4 with M2 polarization activity, further validating known cytokine-process associations. Net formation is strongly associated with GM-CSF signaling^48^.

Taken together, CytED-derived process activities recapitulate known biology and, in conjunction with cytokine signaling analysis, allow for the hypothesization of process-cytokine associations in tumor samples.

### CytED provides a prior over primary and secondary effects of cytokines

Cytokines act in signaling cascades, where perturbation of a cell type with one cytokine can trigger the release of many others, which can then cause production of further cytokines as a secondary effect. In a given perturbation dataset, it is therefore often unclear whether an observed response is the result of a primary cytokine or of its released downstream cytokines (**Fig. 4f**, cf. **Fig. 2f**). We reasoned that we could use CytED to propose a possible disentanglement of such mixed primary and secondary effects. We again use the PBMC 24-hour cytokine stimulation dataset as the experimental design makes secondary effects likely.

Experimentally, IL-12 stimulation of PBMCs results in the expression of 16 cytokines across 6 cell types (28 unique pairs total, **Fig. 4g**). We retrieve all literature connections that match on the input cytokine, cell type, and output cytokine. For IL-12, this yielded only upregulation of IFNγ in natural killer cells as a high confidence connection. Then, we identified all literature-annotated connections for cytokines regulated by IFNγ in any of the cell types present in the dataset, inferring for example that secondary IFNγ was likely responsible for upregulation of CXCL9, CXCL10, and CXCL11 in monocytes (**Fig. 4h**). IFNγ literature annotations overall explained 13 out of 28 of the observed cytokine responses, while IL-12 stimulation itself only explained two. The absence of literature annotations for some of the observed interactions mark potentially novel or underdiscussed findings.

Next, we applied this approach to the signaling events induced by IL-1, which could be explained by a complicated mixture of IL-1 itself, IFNγ released by natural killer cells, and TNFα, IL-6, and CXCL10 released by myeloid cells (**Fig. 4i**). IFNγ broadly regulates secondary cytokines across immune cell types, while the secondary effect of TNFα is more prominent in myeloid cells. To further support these conclusions, we cross-referenced these analyses with CytED-based cytokine activity inference (cf. **Fig. 4a**), subsetting to regulated cytokines with expressed receptors (**Fig. 4j**). For IL-12 stimulation, there is strong IFNγ-signaling across all cell types except NK CD56hi, where IL-12 itself shows the strongest signaling activity. For IL-1 stimulation, IFNγ signaling tends to dominate, consistent with its stronger explanatory power in cytokine signaling connections, with contributions of TNFα and IL-1 in CD14 monocytes, and CXCL10 in NK CD56hi.

### Designing a maximally informative combinatorial cytokine perturbation screen

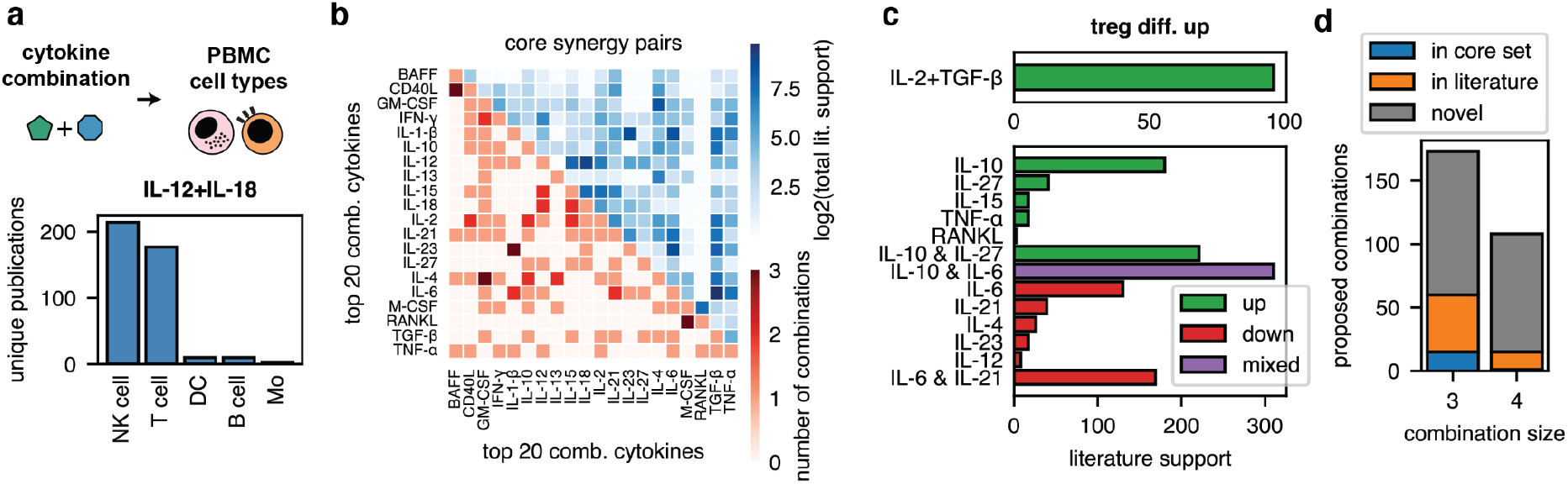
CytED supports the design of cytokine perturbation screens. **a**, IL-12+IL-18 is the most common combination for natural killer cells. **b**, Number of chosen core combinations per cytokine pair and their total literature support (sum over unique publications for each combination). **c**, Literature counts supporting Treg differentiation by IL-2+TGFβ, IL-2 and TGFβ individually, and other cytokines that either support or oppose Treg differentiation. Combining these other cytokines with IL-2+TGFβ yields new informative combinations. **d**, Coverage of proposed new combinations derived by a process as described in (j) by existing core combinations and the literature at large. ‘Novel’ combinations are not annotated even once in CytED.

CytED can guide the design of future experiments in addition to helping analyze existing ones. Here, we provide an example in the context of combinatorial cytokine screening. Cytokines act in concert to generate effects: Sepsis is recapitulated by a combinatorial treatment of mice with TNF plus IL-18, IFNγ or IL-1β but not any individual cytokine^49^. IL-2 and TGFβ synergize to induce stable *FOXP3* expression and peripheral Treg differentiation from naïve CD4^+^ T cells^50^. However, due to the exponential growth in potential conditions, comprehensive and informative high-throughput screens of combinations are hard to design effectively, especially for higher order combinations. Such a screen should include a well-informed selection of known biologically relevant combinations at meaningful concentrations, requiring a comprehensive engagement with prior literature. At the same time, it should include the most promising, currently understudied combinations. We here used CytED to design such a screen, targeting a budget of around 400 combinations.

We first derived a selection of known core literature combinations of two or more cytokines. With PBMCs as a target system, we subset to statements about B cells, T cells, NK cells, monocytes, or dendritic cells. Counting how often each cytokine occurred in combination, IL-12+IL-18 was the most commonly mentioned combination for NK cells, leading to particularly strong release of IFNγ (**Fig. 5a**). We further found that IL-4 and IL-21 are often used in combinations for B cells, while TGFβ- and IL-6-containing combinations are commonly discussed for T cells (**Fig. S18a-b**). We generated a list of just 20 cytokines that covered between 57% (dendritic cells) and 86% (NK cells) of all cytokine combination entries in CytED. Balanced aggregation of literature combinations across these 20 cytokines and 5 cell types yields a core list of well-established combinations (**Fig. 5b**, Methods, **Table S7**), resulting in 75 two, 22 three, and 2 four cytokine combinations, e.g., IL-1β+IL-23+IL-6+TGFβ for particularly potent Th17 differentiation. The particular concentration used for a combination is critical for its observed effects. Using CytED metadata, we retrieved distributions of concentrations used in *in vitro* experiments for the 20 individual cytokines (**Fig. S18c**) and also particular concentrations used for the chosen combinations (**Fig. S18d**). For example, IL-6+TGFβ is most commonly used at concentrations of 20 and 2 ng/ml, respectively.

To derive informative novel combinations, we first observe that these prominent literature combinations often impact on cell differentiation processes. For example, IL-6+TGFβ has Th17 differentiation as its most prominent process (**Fig. S19a**). The individual cytokines also impact on the same processes, albeit much less prominently (**Fig. 5c**). Treg differentiation is the most common process for IL-2+TGFβ, while for IL-2 itself, it’s not in the top 10 most common statements. One way of choosing potentially informative understudied combinations of size *n+m* is therefore to combine *m* cytokines that individually impact positively or negatively on a chosen set of processes, with existing literature combinations of size *n* that impact on that process. For Treg differentiation, this yields, for example, IL-6 or IL-10 plus IL-2+TGFβ, which could clarify the impact of IL-6 on this process (cf. **Fig. 2e**). It also rediscovers IL-1β+IL-23+IL-6+TGFβ for Th17 differentiation (**Fig. S19b**). Applying this scheme to 13 processes and genes yields 173 triple combinations and 108 quadruple combinations (**Fig. S19c, Table S7**). The 173 triple combinations cover 68% of chosen canonical literature triple combinations compared to an expected value of 2.7% for randomly adding another cytokine to the existing two cytokine combinations, suggesting that this process prioritizes biologically plausible combinations. For the quadruple cytokine combinations, around 14% have at least some minimal support in CytED, as compared to an expected value of <1% for random sampling on top of existing combinations. The vast majority of picked quadruple combinations do not occur in CytED at all, presumably because quadruple combinations have largely not previously been studied (**Fig. 5d**). In total, the process yields 384 testable conditions (20 individual cytokines, 99 established literature combinations, and 265 novel combinations).

## Discussion

Here we present CytED, a database of cytokine-cell type-effect statements generated by parsing and multi-step processing of 110,000 full publications using an LLM reasoning model. By converting free-text claims into standardized entries anchored to specific publication excerpts, CytED turns a diffuse literature into an operational resource, allowing us to quantify consensus, similarity, and context dependence in the parsed corpus. CytED thereby enables integrated analyses of literature claims about cytokine behavior and serves as an entry point for highly targeted LLM-powered literature analysis. We provide code and a web interface for CytED, which makes it a searchable index of the parsed immunology literature.

CytED is structured to directly interface with newly provided experimental datasets, serving to support many aspects of a typical analysis workflow. Initially, CytED globally quantifies agreement with observed literature responses, subset on various metadata such as time points or species, if desired, to provide an overview over experimental results. CytED also yields key genes that dominate these agreement numbers and can be individually investigated. Because these key genes retain links to original publications, their biological implications for the experiment can also be scalably and traceably analyzed by LLMs in the context provided by the primary literature. To further support dataset analysis, we used CytED to generate several analytical tools, namely weighted gene signatures for cytokine and cell process activity inference, and literature priors that helped disentangle experimental results into likely primary and secondary cytokine responses. Finally, we showed how CytED can be used to design a sparse combinatorial cytokine perturbation experiment that balances the inclusion of known and the exploration of novel combinatorial cytokine biology.

There are some limitations to our current work: First, CytED inherits the biases of the literature. Literature support for a given relation reflects attention as much as biology, so well-studied cytokines and canonical contexts dominate. This bias is partly useful: We found that high-frequency statements tend to generalize better and literature counts were surprisingly effective for predicting cytokine signaling activity. Still, biases could also obscure biologically key but less studied connections. Thus, CytED should first and foremost be seen as providing useful priors rather than ground truth. Second, coverage is still incomplete: even at 110,000 publications, CytED represents only a fraction (perhaps ∼10%) of the available cytokine research. Further scaling would strongly improve coverage and robustness, but is difficult due to open-access restrictions. Third, not every complex interaction in immunology fits into CytED’s target format. While this is necessary for computational tractability and partly alleviated by the presence of metadata, complex experimental context remains hard to fully consider. Finally, although modern LLM reasoning models are now sufficiently advanced to enable this type of literature aggregation, we found that they still make mistakes, and require extensive prompt and response format testing and tuning as well as repeated quality controls for acceptable results (cf. **Fig. S2**). They are also still expensive when applied at scale. Future model improvements are certain to reduce both the cost, quality, and ease of construction of CytED-style databases.

Beyond its immediate use in immunology, CytED illustrates a general paradigm for facilitating the interplay of high-throughput screens, established literature, and AI models. Much biological knowledge has been discovered through painstaking individual experiments, but is not present in a form that interfaces well with modern high-throughput datasets or computational reasoning. Converting domain literature into standardized, quality-controlled tables custom designed to fit the structure of particular high-throughput experiments could provide reference knowledge substrates across many areas. With dropping costs for LLM use and an increasing scale and cost for high-throughput screens, it seems increasingly feasible to perform such custom literature aggregations to support the design and evaluation of individual screening campaigns. Further, the goal of many high-throughput perturbation datasets currently being produced is to ultimately feed virtual cell models^51,52^. By compressing the literature into a format analogous to experimental data, CytED-style databases provide a knowledge graph against which learned representations can be regularized, validated, or iteratively refined. Rather than treating decades of accumulated biological insight as external commentary, this could enable predictive models to be trained under explicit alignment with established knowledge.

## Methods

### Database construction

#### Publication Data Source

Papers were bulk downloaded from Europe PMC^26^. XML bulk files were converted to JSON format using scripts from the Semantic Scholar Open Research Corpus^53^ and tagged with citation count information from the Semantic Scholar Academic Graph Dataset^27^. One large advantage of EuropePMC is that it provides articles as structured XML files. This avoids PDF parsing, which is error-prone, generates incorrect unicode formatting, and necessitates chunking strategies that might not reflect the logical organization of scientific publications.

#### Curation and ranking of publications

The bulk downloaded dataset was filtered with PubMed search queries. We manually curated synonyms for cytokines using resources such as EntrezGene^54^ and UniProt^55^, but avoided outright copying of these resources as they contain some unfortunate entries for historical reasons (e.g., IL-21 as a synonym for IL-22). This resulted in the cytokine tables in **Table S1** and **Table S2**. We used these names and synonyms for an exhaustive search via the PubMed API using each as an individual search term and noting which publications were returned for each. This yielded 383889 unique publications that were present in our publication repository. We further filtered and ranked papers based on the following steps:

##### 1. Relevance Scores

Each paper in the filtered dataset was initially processed with Gemini-2.0-Flash, which generated a summary and relevance score (1-10) based on a description of our cytokine effect extraction task (*prompt: summarize_publication*). Papers with a relevance score of 5 or below were discarded. For the remaining papers, relevance scores were min-max normalized before combination with citation-based scores. In the same step, we also classified publications as either original work or reviews.

##### 2. Citation scores

Publications are binned by age into intervals of 3 months, then within-bin citation percentile is calculated based on citation count information from the Semantic Scholar Academic Graph Dataset^27^. This adjusts for the fact that older papers have had more time to accumulate citations.

##### 3. Overall paper score

We calculated an overall paper score as the average of normalized relevance score and age-adjusted citation percentile.

##### 4. Publication selection

First, we indexed each paper by the cytokines mentioned, i.e., which PubMed queries found this particular paper for a given cytokine. Publication selection was then performed with a two-phase greedy procedure. In the first phase, we ensured that each cytokine was represented at least once by iteratively selecting papers for the currently rarest uncovered cytokine, preferring papers that simultaneously covered multiple still-uncovered cytokines, with the overall paper score used as a tiebreaker. In the second phase, we expanded the corpus while maintaining balanced cytokine representation. Each cytokine was assigned a target proportion based on its priority (**Table S1**), with higher-priority cytokines receiving greater target weight. At each step, each candidate paper was rescored according to both its overall paper score and how much it improved representation of undercovered cytokines relative to these target proportions.

We selected 100,000 primary papers and 10,000 review papers.

#### Preprocessing of Publication Texts

We converted full text papers from XML to JSON using the doc2json package^53^ (https://github.com/allenai/s2orc-doc2json), providing parseable section headers and inline reference statements. To avoid overly long context-lengths and increase recall when passing texts to LLMs for parsing, publications were split by section header. The figure captions of all mentioned figures and the full table for all mentioned tables in a section were appended exactly once to each publication section.

We generated context for each paper containing a high level summary of the content and methods as well as information on cell type markers, used concentrations, experimental time points, and abbreviations using Gemini-2.5-Flash. Given that standalone paper sections sometimes require an understanding of the paper’s overall goals and methods, these concise summaries were prepended to paper sections to provide LLMs with needed context (*prompt: summarize_publication*).

#### Initial extraction of cytokine effects

We prompted Deepseek-R1-0528 via its API with each of the contextualized paper sections and an instruction to extract statements from the paper that describe cytokine effects on cell types (*prompt: extract_cytokine_effect*)

The result was parsed as a list of JSON objects, each containing the following fields:

*key sentences, causality description, causality type, cytokine name, cell type, cytokine effect, experimental system, experimental perturbation, experimental readout, experimental time point, experimental concentration, necessary condition, citation ID, confidence score*

To minimize data loss, responses that contained malformed JSON or exceeded context length were retried. The model was prompted to separate different entries within one column by a semicolon, e.g., if multiple cytokines, cell types, or experimental systems apply to a single entry. To enforce consistency in experimental system descriptions, LLMs were prompted to format the *experimental system* field for each entry as <species>:<system type>:<details>, where *species* indicates the animal model used (if applicable), *system type* is in vivo/in vitro/in silico/other, and *details* is any additional free form text describing the system. The LLM was instructed to substitute ‘unknown’ for any component that cannot be determined. Entries with a *confidence score* below 7 were removed from the final table. We did not retain experimental metadata information for cited works as we found larger hallucination and guessing by the LLM along with less useful detail.

#### Statement deduplication for chunking strategy comparison

Within each source document and chunking strategy, extracted statements were embedded with text-embedding-3-small and clustered by connected components of a cosine-similarity graph (threshold 0.95), thereby grouping transitively similar statements. Single-statement groups were kept as is. Multi-statement groups were then deduplicated with an LLM (gemini-2.5-flash) instructed to return only a subset of the input statements representing unique cytokine-cell type-effect assertions, without introducing new statements (*prompt: deduplicate_statements*). The resulting count of retained statements was used to compare chunking strategies.

#### Initial quality control

Rows extracted by LLMs were filtered based on grades assigned by an LLM judge. To implement this, we prompted the same model (Deepseek-R1-0528) with the original text passage, the list of LLM-extracted contents, and detailed instructions on how to assign grades to individual responses. Each response received a separate grade for *cytokine name, cell type, basic interaction* (effect description), *causality type, citation type, citation id, experimental readout, experimental perturbation*, and *necessary condition*. The scale of grades for each category generally included ‘correct’, ‘approximate’, and ‘incorrect’, but additional category-specific labels to identify error modes at a more nuanced level (*prompt: qc_initial_extraction*). Entries with ‘incorrect’ in *cytokine name, cell type*, or *basic interaction* were removed from the final table. Failed entries in other columns were converted to ‘unknown’. We note that this quality control step used mapped cytokine and cell type names (see below) and therefore also assessed mapping quality of these columns relative to the original text passage, i.e., whether the mapping process destroys the validity of the triple.

#### Postprocessing cytokine effects

To facilitate usability in downstream analysis workflows, we extracted regulated genes, pathways, and cell processes from the free text *cytokine effect* descriptions extracted in the first step. To generate this, we again prompted Deepseek-R1-0528 with entries (limited to fields *cytokine name, cell type, cytokine effect, experimental system, causality description, key sentences*) from the initial extraction and an instruction to extract a list of JSON objects with the following fields (*prompt: process_cytokine_effect*):

*cytokine effect, regulated genes, gene response type, regulated proteins, protein response type, regulated pathways, pathway response type, regulated cell processes, cell process response type*

Protein-level responses were not further processed for this publication but were necessary to prevent the LLM from entering purely protein-level responses into the *regulated genes* column.

#### Cytokine effect postprocessing quality control

To evaluate the quality of cytokine effect postprocessing, we used an LLM judge (DeepSeek-V3.2) to determine if the regulated genes, cell processes, and pathways extracted are consistent with the original cytokine effect described (*prompt: qc_effect_processing*). Each category received an overall grade (‘correct’, ‘incorrect’, or ‘not mentioned’) and also a grade for the direction.

#### Evaluation of gene symbol extraction

Postprocessing free text cytokine effects is moderately complex, requiring that the LLM correctly infers whether or not a gene is implied, outputs the valid gene symbol, and determines whether the effect described refers to an mRNA expression change, a post-translational protein modification, or a modified pathway. For example ‘upregulation of cell adhesion molecules’ in epithelial cells might reasonably imply *ICAM1, VCAM1*, or *SELE*, though not obviously so. To assay the ability of different LLMs to perform this task, we manually reformulated 52 rows, generating 118 output rows containing 116 implied mRNA-level changes. The manual reformulation was performed in such a way that all reasonably inferrable mRNA changes (by our judgement) were exhaustively annotated, implying that additional annotations introduced by an LLM would be erroneous (**Table S8**).

#### Ontology Mapping

To map free text cell types and pathway names to standard ontology terms, we used the following algorithm:

1. Input terms are rewritten using an LLM (gemini-2.5-flash) to partially standardize the syntax. Specific characteristics prompted include the removal of extraneous qualifiers, standardization of separators when multiple cell types are listed, and expansion of known abbreviations. For the cell type mapping task specifically, the LLM was also prompted to classify each term as ‘body’, ‘cell line’, ‘cancer’, or ‘other’ (*prompt: process_cell_type*). For example, *monocytes from rf patient (m0 phenotype)* would be rewritten as *m0 monocyte* and labeled as *body*.
2. Each of the rewritten terms was then mapped to a target ontology term. To implement this, we first generate dense semantic embedding vectors (d=1536) using OpenAI’s text-embedding-3-small model for all terms (and their associated synonyms) in the ontology and calculate the cosine similarity of each to the cell type being mapped. We also calculate a syntactic similarity score based on sparse 3-character level TF-IDF vectors. To capture both semantic and syntactic similarity, we use a linear combination of the above similarity metrics, specifically:

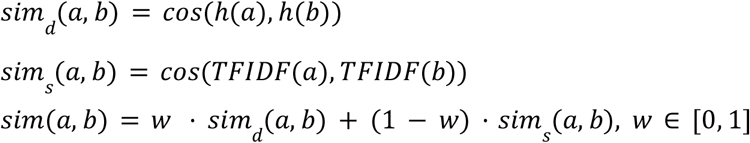

We calculate *sim*(*a, b*) at *w* = 0, 0. 25, 0. 5, 0. 75, 1. 0, take the top 10 scoring terms at each weight, and filter the candidate term list to those with a similarity score of at least 0.6. We then prompt Gemini-2.5-Flash to select the most fitting target cell type among these candidates, or report that the term failed to map (*prompt: choose_cell_type*).

#### Cell Type Standardization

Body cells were mapped to a filtered version of Cell Ontology^28^ (2916 IDs, 80.7% of mapped entries), cell lines were mapped to a filtered version of Cellosaurus^30^ (141821 IDs, 11.1% of mapped entries), cancer cell types were mapped to a filtered version of the NCI Thesaurus^29^ (2292 IDs, 8.2% of mapped entries), and ‘other’ entries were discarded. Ontologies were downloaded as OBO files and processed using the Obonet python package. For Cell Ontology, we removed redundant species-qualified labels when an unqualified term was present (for example, ‘, human’), excluded malignant terms, and then collapsed remaining species-qualified names to their generic form. A few terms were added when the detail provided by Cell Ontology was insufficient. For NCI Thesaurus, we restricted the ontology to descendants of the neoplasm node and to terms annotated as belonging to a neoplastic process, then filtered nodes to retain general cancer-type terms while excluding overly specific nodes, including terms stratified by species, treatability, age, or stage; parent nodes required to preserve connectivity in the retained hierarchy were kept. Cellosaurus terms were lightly normalized by removing parenthetical qualifiers and duplicate aliases.

For Cell Ontology and NCI Thesaurus, we used the procedure described in Ontology Mapping to convert the cell types extracted from papers to their closest ontology equivalent. For Cellosaurus, all terms are standardized by lowercasing and removal of any non-alphanumeric characters and mapped using 3-character TF-IDF similarity only. To evaluate the quality of cell type mapping, we compared ontology CURIEs mapped by our algorithm to 440 manually annotated entries (**Table S9**).

#### Cytokine Standardization

Raw cytokine names were standardized using a multi-step normalization and synonym-mapping pipeline. Names were converted to lowercase, unicode-normalized, and stripped of parenthetical annotations, extraneous characters, and non-informative descriptors (e.g., ‘recombinant’). Greek letters and whitespace were harmonized to a dash-separated format. Processed names were then mapped to canonical cytokine identifiers using a curated synonym table derived from manual curation of individual cytokines and cytokine families. The synonym dictionary was expanded computationally to include systematic variants, including alternative prefixes (e.g., species or recombinant annotations), dash insertion/removal, abbreviation substitutions (e.g., IFN/interferon), and pluralization variants. Compound cytokine annotations separated by delimiters (‘+’ or ‘;’) were parsed and mapped component-wise. Entries that could not be unambiguously mapped were labeled as unmapped and excluded from downstream analyses.

#### Gene Symbol Standardization

Cytokine effect entries with non-empty *regulated genes* reported were filtered to those recognized as a valid gene symbol. We collected a lexicon of valid gene symbols by combining the following sources:

1. All gene names from the *human, mouse, rat, pig, cow, chicken, dog, rhesus, chimpanzee, frog, zebrafish, horse, sheep*, and *rabbit* Ensembl Gene datasets (accessed via the pybiomart python package).
2. Known gene symbol versions of all cytokine names in our manually curated vocabulary (e.g. IL-12 to IL12A, IL12B; TGF-β1 to TGFB1). Matches to a cytokine’s non-gene name were converted to the gene symbol version.
3. Human and mouse gene symbols from the NCBI HomoloGene and Ensembl Gene databases (accessed via the mygene python package). Matches to registered aliases of a gene symbol were mapped to their canonical versions.

Only 0.96% of LLM-extracted gene symbols did not match any in the lexicon. Entries containing invalid gene symbols were filtered out from the knowledge base.

#### Pathway Standardization

Pathways were mapped to terms in the NCBO Pathway Ontology^31^, which organizes biological pathway names into a tree hierarchy with a maximum depth of 9 layers. To select a relevant subset of terms, we used terms in the signaling (PW:0000003) and regulatory (PW:0000004) categories. To get the desired level of granularity, we kept terms 2-6 layers deep in the ontology tree. However, for terms within the cytokine & chemokine signaling (PW:0000577), signaling pathway pertinent to immunity (PW:0000818), and immune response pathway (PW:0000023), all terms were kept regardless of depth. This resulted in a total of 800 canonical terms, and an extended vocabulary of 1649 terms that includes synonyms listed with each entry in the ontology. LLM-extracted pathway names were mapped to ontology terms using the same process described in Ontology Mapping (Methods) with different prompts (*prompt: postprocess_pathways, map_pathways*). Of the 169,630 pathway names extracted in CytED, 79.0% were mapped to a term in the Pathway Ontology (416 unique concepts).

To address gaps in the coverage of ontology terms, we prompted Gemini 2.5 Flash to judge the validity of each mapped pathway name assigned compared to its original counterpart (*prompt: check_pathway*). Specifically, each pathway entry was deemed ‘correct’, ‘incorrect’, or ‘approximate’. If judged ‘incorrect’, the LLM was also instructed to output an alternative name that is more fitting. After this, a final round of manual corrections was applied to standardize the syntax of commonly occurring alternative pathway names that didn’t map to a Pathway Ontology term in the first round (primarily STAT and SMAD related concepts).

#### Cell Process Standardization

Given the relatively small number of unique cell process terms extracted without any preprocessing, we curated our own set of cell process categories. As a starting point, we ordered the ‘cell process’ terms in CytED by frequency and took the minimal set that accounted for 50% of entries (18 unique terms). To widen the vocabulary, we used an LLM (Gemini-2.5-Flash) to extract key core terms from the remaining 50% of entries. Specifically, we sampled 20 batches of 50 cell process terms each, and prompted Gemini-2.5-Flash to output lists of non-redundant terms/categories that these processes can be grouped into (*prompt: categorize_cell_process*). We favored more frequently-occurring terms while sampling, assigning sampling weights based on term’s rank when ordered by frequency in CytED. The results for each batch were merged. Using this initial set as guidance, we then manually curated 263 cell process terms using expert judgement (**Table S4**). Using this vocabulary, we mapped the original set of cell process descriptions to vocabulary terms using strategy described in Ontology Mapping (*prompt: map_cell_process*). After mapping, we added a consistency check using an LLM (Gemini-2.5-Flash) to detect either mismatched terms or those with flipped polarity (e.g., ‘bone resorption’ mapping to ‘bone formation’, *prompt: process_consistency_check*).

#### Experimental Metadata Standardization

For the *experimental system* descriptions extracted in the initial cytokine effect extraction step, we used Gemini 2.5 Flash to repair a handful of entries that were either ill-formatted or contained long overly-specific descriptors (*prompt: parse_experimental_system*). If an entry could not be inferred from its ill-formatted version, it was deemed ‘unknown’. For the remaining entries, the *experimental system* field was split into separate metadata fields *species, experimental system type, experimental system details*.

To facilitate downstream analysis of the *experimental concentration* fields, we used a variety of regex filters to extract numbers and units and to eliminate entries that did not report quantitative values (e.g. ‘not specified’, ‘overexpression’). To reduce ambiguity of unexpected parenthetical statements or malformed text, we prompted Gemini-2.5-Flash to reformat each set of concentration values as a list of records formatted as <entity>:<value> <unit>, where *entity* is the cytokine treatment, *value* the numeric quantity, and *units* the measurement unit (e.g. ng/mL, μg, etc.). The list was followed by an annotation that identifies which items were part of the same treatment group, if any (*prompt: parse_concentrations*).

*To map the experimental time point field*, we parsed free-text time annotations using regular expressions that captured common formats including explicit value-unit pairs (for example, 24 h, 2 days, 30 min), range expressions (for example, 24–48 h or 24 to 48 h), and developmental shorthand such as Day X. Extracted units were normalized to a standard set (minutes, hours, days, weeks, months, years) and converted to hours. Entries that were missing, unknown, or did not contain interpretable quantitative time information were excluded. For downstream use, parsed values were serialized into a structured representation that preserved both the original text and the processed hour values.

Conditions described in the *experimental perturbation* field were postprocessed using Gemini-2.5-Flash (*prompt: categorize_experimental_perturbations*) into JSON objects with fields *category* (one of Compound treatment, Genetic manipulation, Physical stimulus, Environmental stress, Microbial/Viral, or Complex/Contextual), *entity* (a list of entities used in the perturbation), *direction* (the intended effect of the perturbation, e.g. ‘increase’, ‘decrease’, ‘modulate’, or ‘unknown’), and *additional info* (any extra descriptors, e.g. dosage, method of administration, etc.). Entries that failed the first round of post-processing were rewritten (*prompt: rewrite_perturbation*), then underwent another round of attempted categorization.

Values in the *experimental readout* field and had fewer structural constraints and were processed and mapped following the same procedure as cell processes with adjusted prompts (*prompt: categorize_experimental_readout, map_experimental_readout*).

#### Human quality control

To assess the effect of QC on the core triple (cytokine-cell type-cytokine effect), we used 1000 pre- and 2000 post-QC table rows, both after all mapping and post-processing steps, were evaluated for the presence of clear errors using GPT-5.2 as a judge by comparing to the original text passage (*prompt: qc_model_comparison*). The prompt is structured to make it more likely to only detect true errors, while the original qc prompt is structured to overpredict errors to increase precision. We then extracted all entries labeled as ‘incorrect’, merged entries from the pre- and post-QC table, and shuffled them to obscure their origin. These shuffled entries were then manually re-evaluated, allowing us to estimate relative error rates pre- and post-QC.

To assess error rates in the final, quality controlled table, we sampled 300 text passage ids and retrieved all associated rows in CytED for a total of 1348 rows. We re-ran the initial QC (*prompt: qc_initial_extraction*) using GPT-5 as a judge as an initial estimate of the error rate. To contextualize GPT-5 ratings, we then subsampled the chunks to those with mixed ratings in their rows (at least 1 ‘correct’, at least one ‘extrapolated’ or ‘incorrect’), yielding 134 rows for manual annotation. The goal of the subsampling is to reduce the number of manually annotated rows necessary to estimate an upper limit to the error rate under the assumption that an entry labeled as incorrect by GPT-5 is more likely to be genuinely incorrect. The subsampled rows were manually evaluated against their original text passage.

### CytED analysis and aggregation

#### Cell type and cytokine coarse-graining

To coarse-grain cytokines, we mapped cytokine names to the ‘summary_cytokine’ column in **Table S1**. To aggregate literature statements at different cell type granularities, fine-grained Cell Ontology identifiers in CytED were either mapped to broader categories using upward traversal of the Cell Ontology, or a higher-level node was distributed across more specific cell types: Each statement for ‘T cell’ was also assigned to ‘CD4 T cell’ and ‘CD8 T cell’, each statement for ‘monocyte’ was also assigned to ‘CD14-positive monocyte’ and ‘non-classical monocyte’, and each statement for natural killer cell was also assigned to ‘CD16-negative, CD56-bright natural killer cell’ and ‘CD16-positive, CD56-dim natural killer cell’. For lower-level nodes, statements were upconverted to the nearest cell type annotated in the experiment.

#### Deriving literature statement counts

Cytokine-cell type-entity-direction count tables are calculated by aggregation across CytED based on several parameters. First, cytokines and cell types are either taken as is or coarse-grained (see above). Second, the *causality type* field can be constrained to ‘direct’ or ignored. Third, directions can be aggregated by subtracting the larger direction count from the smaller direction count and retaining only the larger count, or neither if they were equal. Fourth, CytED can be further constrained based on metadata columns such as species or citation classification. For all count tables, we dropped duplicates in cytokine-cell type-entity-direction within a publication. For the baseline counts tables, we did not perform cytokine, cell type or direction aggregation or causality filtering.

Confidence labels (low, medium, high) for a given set of cytokine-cell type-entity-direction counts are assigned separately for each cytokine based on the count percentiles across that cytokine’s cell type-entity pairs. The thresholds for medium and high confidence correspond to the 80th and 95th percentiles of weights, subject to absolute minima of 2 and 4 counts, respectively.

#### Cytokine-cytokine regulation graph construction

To find cytokine-cytokine regulatory relationships, we created a cytokine-cell type-gene-direction count table as described above, with coarse-grained cytokines and causality filtering. Duplicates within a publication were dropped. We then identified cytokines whose known subunit genes (‘gene’ in **Table S1**) appear as regulated targets in the literature. A regulation entry was retained only when all subunits of the cytokine were reported as regulated by the same input cytokine in the same cell type and direction. The weight of each cytokine-cytokine regulatory statement was taken as the minimum count across component genes.

#### Higher-order effect analysis

To infer higher-order effects, we first calculated and merged non-coarse-grained count tables for genes and cell processes with constrained causality and direction aggregation. Then, we combined the cytokine-cytokine regulation graph and this combined table by merging them on the output cytokine of the regulation graph and the input cytokine of the combined table, e.g., every effect for IL-6 in the latter is associated with each cytokine-cell type combination containing regulation statements for IL-6 in the former. The graph is subset to a particular cytokines and cell types of interest at every step, e.g., anything with an impact on m2 polarization as the final step. For this subset graph, the support of the two steps is calculated as log2(count + 1) divided by the maximum value for the subset graph. The overall support is calculated as log2(count_1 + 1) + log2(count_2 +1) divided by the maximum value for the subset graph.

#### Similarity and UMAP embedding analysis

To analyze the similarity between cytokines, we calculated and merged non-coarse-grained count tables for genes and cell processes with constrained causality and direction aggregation. Cell processes were filtered to remove low-information entries (e.g., ‘gene expression’, see ‘specificity’ in **Table S4**). Downregulation counts were given a negative sign. Each cell type-entity pair was then treated as a feature. For across-cell type feature vectors, we first filtered to features with at least 10 counts for one cytokine and at least 3 counts for at least 3 cytokines. For within-cell type feature vectors, we first filtered to features with at least 5 counts for one cytokine. In both cases, we then filtered cytokines to those with at least 100 total effective counts and excluded cytokine combinations. The resulting feature vectors were normalized so that their absolute values sum to 1.

To create UMAP embeddings, we performed truncated singular value decomposition on the feature vectors, ran UMAP using the python umap package^56^ with n_neighbors=30, min_dist=0.1, and a cosine metric.

#### LLM-based literature traversal for hypothesis formation

For a particular relation of interest such as the influence of IL-6 on *FOXP3* expression in regulatory T cells, we subset CytED to all entries matching this triplet. We split the resulting rows according to whether they were cited statements or original research findings. We then retrieved all associated literature chunks for original statements. For each such literature chunk, we provided Gemini-2.5-Flash with the consensus literature (cited) observation for the relationship, then asked it to summarize the text chunk in light of three points, namely what the text chunk reports regarding the regulatory relationship, what its experimental conditions and subtleties are, and how this might inform any potential disagreement (*prompt: chunk_analysis_literature*). We then provided these summaries to GPT-5.2 and asked it to analyze them to provide an overall analysis of the relationship of the literature consensus to experimental findings, as well as brief summaries, key insights, and potential testable hypotheses (*prompt: overall_analysis_literature*).

### Integration of CytED with experimental data

#### Preprocessing of experimental data

To preprocess the human PBMC dataset for comparison, we retrieved response magnitudes from the original publication and filtered cytokines to those that caused a strong response in at least one cell type (B cell, CD4 T cell, CD8 T cell, Treg, NK CD56hi, NK CD56low, cDC, CD14 Mono, CD16 Mono). Cytokines were coarse-grained as described above. When multiple cytokines had the same coarse-grained name, we retained the one with a larger mean response magnitude across cell types (e.g., IL-1-β was kept for the coarse-grained cytokine IL-1). We used provided log2FC values. To estimate an error of the provided log2FCs values, we calculated a standard error using the F-statistic as SE = abs(log2FC)/sqrt(F). From this we calculated log2FC_upper = log2FC + 0.5*SE and log2FC_lower = log2FC - 0.5*SE. The log2FC data was then coarse grained as either upregulated, downregulated, unchanged, or uncertain depending on whether both were above 0.5, below -0.5, in between -0.5 and 0.5, or distributed across these boundaries. Uncertain values were discarded.

To process the in vivo mouse perturbation data, pseudobulk expression profiles were generated separately for each mouse cell type from the original single-cell dataset. First, cells were split by annotated cell type, and within each cell type, only genes detected in more than 5 percent of cells were retained. For each cell type, counts were summed across cells belonging to the same biological replicate within each cytokine condition, yielding one pseudobulk profile per replicate and stimulation condition. Pseudobulk samples supported by 10 or fewer cells were excluded. Genes were filtered per cell type to those with a mean raw pseudobulk expression across replicates greater than 10 counts in at least one cytokine condition. Analyses were further restricted to conditions represented by at least two replicate pseudobulk samples, and each cell type was required to have at least two PBS control replicates. For each cell type, differential expression was then tested across cytokine conditions using edgeR^57^ via the pertpy interface^58^ with a design matrix modeling cytokine treatment and with PBS as the reference condition. Log2FC coarse-graining was then performed as described above for human PBMC data, except that we used boundaries of 1 and -1 to account for higher noise levels.

### LLM-based literature traversal for the interpretation of key experimental genes

To interpret experimental results in the regulation of key genes in the context of the primary literature supporting CytED statements, we implemented a two-stage LLM-based analysis. For a selected cytokine and cell type combination, we identified the top most common literature-supported statements in the integrated CytED-experimental data table (e.g., all statements with at least 5 mentions in the literature). The consensus literature direction was defined based on up- and downregulation counts, with genes with no majority (neither direction exceeding twice the other) being labeled as ambiguous.

We retrieved the full text of all literature passages discussing these key genes in the context of the cytokine and cell type. Each selected chunk was analyzed with GPT-5-mini (medium thinking) in a single query. The system prompt described the experimental setup (cell type, cytokine, stimulation duration, readout modality) and provided the list of key genes with their reference log2FC values, consensus literature directions, and agreement classifications. The model was instructed to (*prompt: chunk_analysis_key_genes*): (1) summarize the experimental conditions and gene regulation described in the chunk (*summary_of_text_chunk*); (2) provide per-gene summaries of cell state implications (*gene_summaries*); and (3) discuss what agreements and disagreements between the literature and the reference experiment imply about the cell state (*implications*).

To integrate the information and synthesize an overall insight into the implications of the experimental results, the per-chunk summaries were then passed together to a second, more capable language model (GPT-5.2 with long thinking, *prompt: overall_analysis_key_genes*) in a single synthesis query. The model was provided the full list of key genes with their experimental context and was instructed to (1) synthesize the individual chunk analyses into a coherent interpretation of the cell state under cytokine stimulation, with explicit reference to specific genes and text chunks (*analysis*), and (2) a concise summary of the analysis (*summary*).

### Cytokine signaling inference

#### Cytokine gene set construction

Pathway gene sets were constructed by manually mapping cytokine names to core transcription factors (see **Table S5**). Weighted gene sets for transcription factors were retrieved from CollecTRI^59^ via the decoupler library^42^ and filtered to those with ‘regulon’ or ‘PMID’ for ‘sign decision’. For each cytokine, gene targets and weights were appended for all core transcription factors. When a single gene target was present in the nets for multiple transcription factors, we took the mean of the weight. CytoSig^11^ gene sets were downloaded from https://github.com/data2intelligence/CytoSig/tree/master/CytoSig/signature.centroid.expand. Cytokine names were mapped to the CytED nomenclature.

CytED gene sets were constructed by aggregating literature data. For each cytokine-gene pair, up- and downregulation counts were tallied across the aggregated literature. Weights were computed as the net regulatory count (upregulation count minus downregulation count), then log2-transformed with sign preservation: weight = sign(net count) * log2(|net count| + 1). Weights within each cytokine group were then normalized to the maximum absolute weight in that group. Gene sets were filtered to cytokines with a minimum of 10 target genes and at least one gene with a net count of 8.

Human CytoSig and pathway-based gene set targets were translated to mouse orthologs using Ensembl BioMart (pybiomart), retaining only 1:1 orthologs with a non-null human gene name. CytED mouse gene sets were derived directly from the mouse gene columns of the CytED database and required no translation.

#### Inference procedure and performance evaluation

For each cell type in a dataset, the subset of sample rows corresponding to that cell type was extracted from the log2FC matrix. All nets were filtered to the same 41 shared cytokines for a fair comparison. Univariate Linear Model (ULM) was applied using dc.mt.ulm(), returning per-sample cytokine activity scores and adjusted p-values. For each sample, the inferred activity score of the true stimulating cytokine was ranked among all scored cytokines. Ranks exceeding 10 were capped at 10 to avoid comparison of failed inferences.

For CytoSig, we also downloaded and installed the original package from https://github.com/data2intelligence/CytoSig, then ran CytoSig.ridge_significance_test() using default parameters with the input data and filtering steps. Although differences to using decoupler with CytoSig gene sets were generally minor, all CytoSig plots in this publication use the results from the actual CytoSig package rather than decoupler.ulm() to more closely align with the original publication.

#### Used datasets

Inference was benchmarked on three datasets: (i) a subset of the CytoSig database^11^, namely all samples with cell types fibroblast, macrophage, T cell, monocyte, epithelial cell, keratinocyte, endothelial cell, dendritic cell, smooth muscle cell, natural killer cell, B cell, or hepatocyte (essentially subsetting on common primary cells instead of cell lines), (ii) a subset of an in vitro human PBMC cytokine screen^4^, and (iii) a subset of an in vivo mouse cytokine screen^2^. Datasets (i) and (ii) were used as provided by the authors. Dataset (iii) was constructed by calculating log2FCs from pseudobulk data provided by the authors as the publication itself only supplies statistically significant regulatory relationships (see above).

Because (i) partially represents the original training data for CytoSig, we did not include it in the method comparison. For the evaluation datasets (ii) and (iii) were subset based on the original publication to include samples that show strong responses with a good change of being primary responses. For instance, the IL-12 response in monocytes is primarily mediated by secondary IFNγ released by other cell types; because our goal is to infer the direct stimulus rather than downstream paracrine signaling, inclusion of this combination would skew the evaluation. They were also subset for a balanced selection across cytokines and cell types to avoid overemphasizing one specific cytokine or cell type in the evaluation (**Table S6**). Pairwise comparisons between methods were conducted using paired Wilcoxon signed-rank tests applied to the combined human and mouse benchmark ranks.

#### Receptor expression filtering

For secondary effect analysis, inferred cytokine activities were subset to cytokines for which the cognate receptor is expressed. A boolean receptor expression mask was computed from the PBS control expression data (CPM). For each cytokine, receptor expression was defined as the minimum expression across all required subunits (provided in **Table S1**) and the maximum across alternative receptor variants. Cytokine activity scores were discarded for cell types in which receptor cpm were below 8. Cytokines without receptor information were retained by default.

### TCGA cytokine and process activity analysis

To generate process gene sets, we counted co-occurrence of genes and processes deriving from the same origin row in the initial extraction step. When directions matched between a gene and a process statement, counts were incremented by 1, otherwise they were decremented by one. We then filtered to processes with at least 30 total gene counts and 5 associated genes. Gene counts were then log2-transformed with sign preservation: weight = sign(net count) * log2(|net count| + 1) and weight vectors were divided by the maximum of their absolute component value.

TCGA data (RSEM tpm) and metadata (TCGA TARGET GTEX selected phenotypes) were downloaded from # https://xenabrowser.net/datapages/?cohort=TCGA%20TARGET%20GTEx. Samples were filtered to those with at least 50 samples in ‘primary disease or tissue’. For the IL-27 activity analysis, samples were further filtered to those containing the word ‘melanoma’. Genes were filtered to informative genes with at least 10 tpm difference between their maximum and minimum values. To test for the expression of cytotoxicity genes, we took the mean of the log2 tpm values of *CD8A, GZMA, GZMB, GZMH, GZMK, PRF1* and *NKG7*. For enrichment analyses, we added a pseudocount of 1 to tpms, then divided each gene by its median tpm across samples, then log2-transformed the data to generate log2FC values. Cytokine and process activity scores were then calculated with dc.mt.ulm() as described above. To test for process activity in different cancer types, we calculated mean log2FC values and 95% confidence intervals across each disease.

To find associations between cytokine activity and process activity, we downloaded leukocyte fraction estimates^60^ from https://gdc.cancer.gov/about-data/publications/panimmune and removed the component of gene expression associated with leukocyte fraction within each cancer type. Specifically, for each cancer type and each gene, we fit a linear model relating expression to leukocyte fraction across samples and retained the residuals as adjusted fold change values, which we used to recompute cytokine and process activities for each sample. For each cytokine and cellular process, we computed Spearman correlations separately for each cancer type, then summarized these correlations by averaging Fisher z-transformed coefficients and back-transforming to the correlation scale, using the back-transformed mean z plus standard deviation and mean z minus standard deviation as errors.

### Secondary effect inference

We here aimed to identify potential indirect cytokine effects mediated through a secondary cytokine, essentially automating what a researcher would do to identify such relationships based on literature data. We applied a two-step inference procedure, starting from a merged table combining literature-derived cytokine-cell type-cytokine regulation (see **cytokine-cell type-cytokine regulation graph construction**) and experimentally observed cytokine-cell type-cytokine regulation. The two tables are merged and labeled as containing either only literature support, only experimental support, or both.

To derive primary coverage, all experimental observations for a cytokine are collected. Those with a matching literature entry are classified as primary covered relationships. For each such primary covered upregulated cytokine, the full set of downstream regulatory effects attributed to that secondary cytokine in the literature is retrieved. At this point, we apply direction propagation: when the secondary cytokine is predicted to be downregulated in the first step, the signs of its downstream literature effects are flipped, because reduced production is assumed to invert expected downstream consequences. The resulting statement set is intersected with the set of previously unexplained experimental observations to identify novel explanatory coverage. Secondary cytokines that explain no previously unexplained observations are discarded.

### Design of a literature-informed combinatorial cytokine screen

We use a rough cell type mapping (cf. Cell type and cytokine coarse-graining) and subset CytED to statements about B cells, T cells, monocytes, natural killer cells or dendritic cells. Gene and cell effect publication counts were aggregated and merged, then subset to statements about cytokine combinations (i.e., those whose cytokines contain a ‘+’). To pick individual cytokines and their canonical cytokine combinations, we counted publications per cytokine-cell type combination. For each of the five PBMC cell types, we then split up combinations and aggregated counts by individual cytokines, taking the top 7 cytokines with the most counts in combinations per cell type, yielding a total of 20 individual cytokines. Combination counts were then subset to those where all cytokines in a combination were in the picked set of 20. To pick canonical combinations, we iterated over each individual cytokine and PBMC cell type, then took the top 4 most common cytokine combinations in that cell type with at least 2 supporting publications containing the individual cytokine, yielding 99 core literature combinations.

To pick novel combinations, we picked a set of 14 target processes or genes, namely: treg differentiation, th1 differentiation, th2 differentiation, th17 differentiation, CD14, macrophage differentiation, m2 polarization, osteoclast differentiation, immunoglobulin production or secretion, immunoglobulin class switch, plasma cell differentiation, inflammatory cytokine production or secretion, cytotoxic effector function, and cell differentiation. For each process, we found the top 3 literature combinations that influence it. For each of these, we then picked the top 5 individual other cytokines that have the same or the opposite direction of influence on the target process, and filtered them to those with at least two supporting combinations. We then generated five synergistic and five antagonistic combinations by adding them to the original combination. If the initial combination had exactly two cytokines, we additionally took the top two synergistic and the top two antagonistic cytokines, and created one synergistic four cytokine combination, one antagonistic four cytokine combination, and one mixed cytokine combination by adding both the top synergistic and the top antagonistic cytokine. In total, this yielded 265 unique novel combinations.

### Statistical analysis

The statistical tests used are described for each analysis in the corresponding text. When not otherwise mentioned, error bars represent the standard deviation.

## Data availability

Data necessary to supplement the provided code for table construction has been uploaded to Zenodo at https://doi.org/10.5281/zenodo.19020292. The finished CytED table and additional analysis results have been uploaded to Zenodo at https://zenodo.org/records/19468106. CytED is also hosted at https://immune-context.github.io/cyted/.

## Code availability

A package for table construction and reproduction of a subset of figures is available at https://github.com/immune-context/papers2db. The CytED package to perform the literature and experimental data analysis steps presented in this publication (LLM-based exploration, cytokine signaling inference, process signaling inference, secondary effect analysis) is available at https://github.com/immune-context/pyCytED.

## Author contributions

L.O., P.W.K., and G.S. conceived the project. L.O. and M.P. developed the LLM extraction pipeline and packages. R.S. advised and assisted with pipeline and software development. L.O. performed comparisons to experimental data and developed the pyCytED package. M.P. developed the web interface. L.O., P.W.K., and G.S. supervised the project.

## Acknowledgements

We thank Thomas Mayer for assistance with manual evaluation of the database.

## Funding Information

EMBO Postdoctoral Fellowship Program (L.O.)

NIH Award R01GM149631 (G.S.)

NIH Award R33CA28694 (G.S.)

Singapore AI Visiting Professorship AIVP-2024-001 (P.W.K.)

Schmidt Sciences AI2050 Early Career Fellowship (P.W.K.)

