## Supplementary Information for "Transforming the cytokine literature into a resource for experimental analysis and discovery"

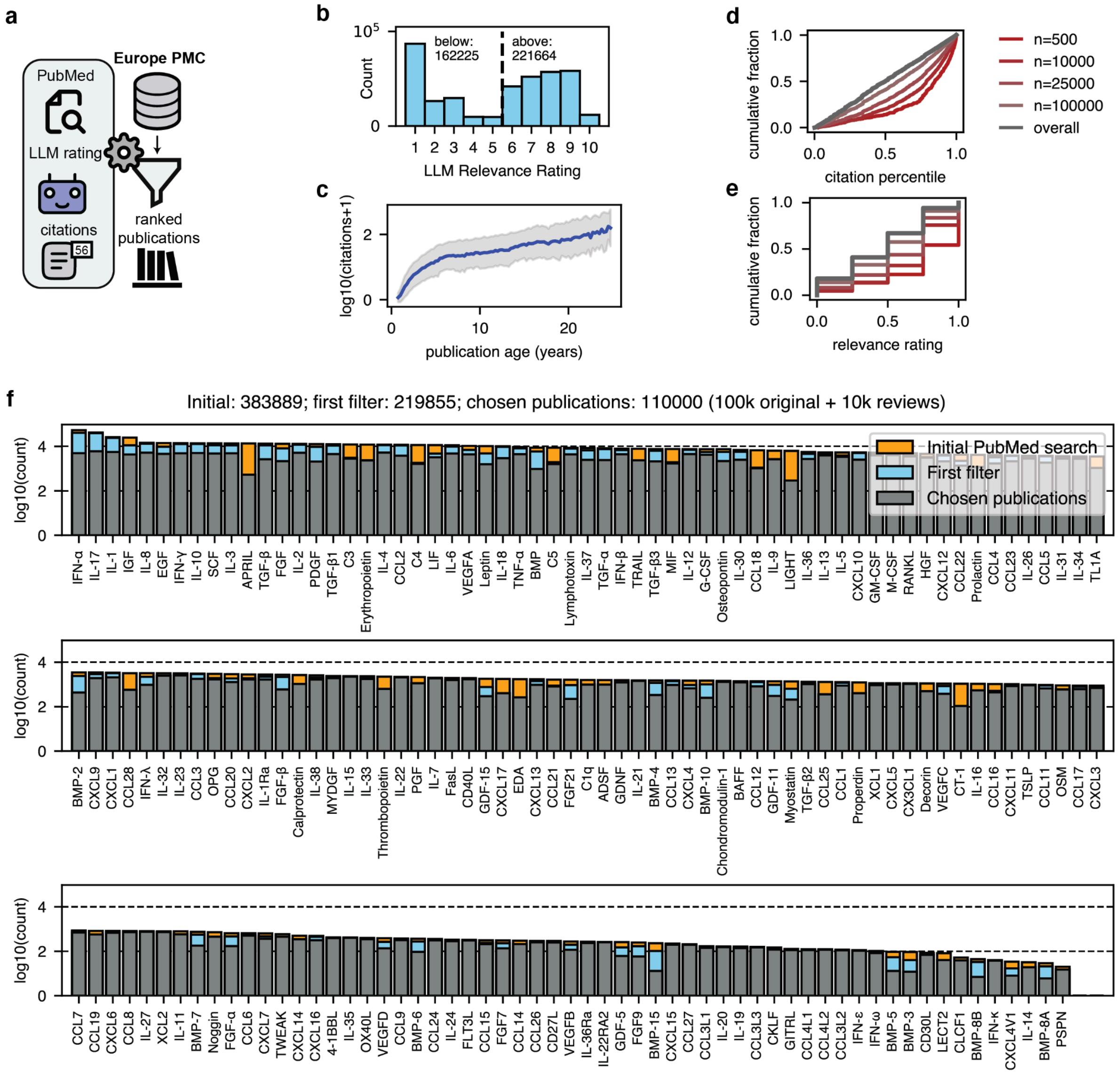


**Fig. S1. Curation of the scientific literature for relevant publications. a,** Open-access articles from Europe PMC are filtered and ranked using a mixture of PubMed Search for cytokine names and their synonyms, LLM-derived ratings from gemini-2.0-flash, and, citation percentiles. **b,** Distribution of relevance scores after PubMed search. Articles with a score less than six are excluded. **c,** Mean and standard deviations of citation counts across three month bins. **d, e,** Distribution of citation percentiles (d) and relevance ratings (e) for the top n papers after the ranking algorithm. **f,** Distribution of publication counts per cytokine (presence determined by a certain publication being a hit for a cytokine or its synonym in the PubMed search) after the initial PubMed Search, the initial relevance score filter (cf. (b)), and after taking the top 110k publications.

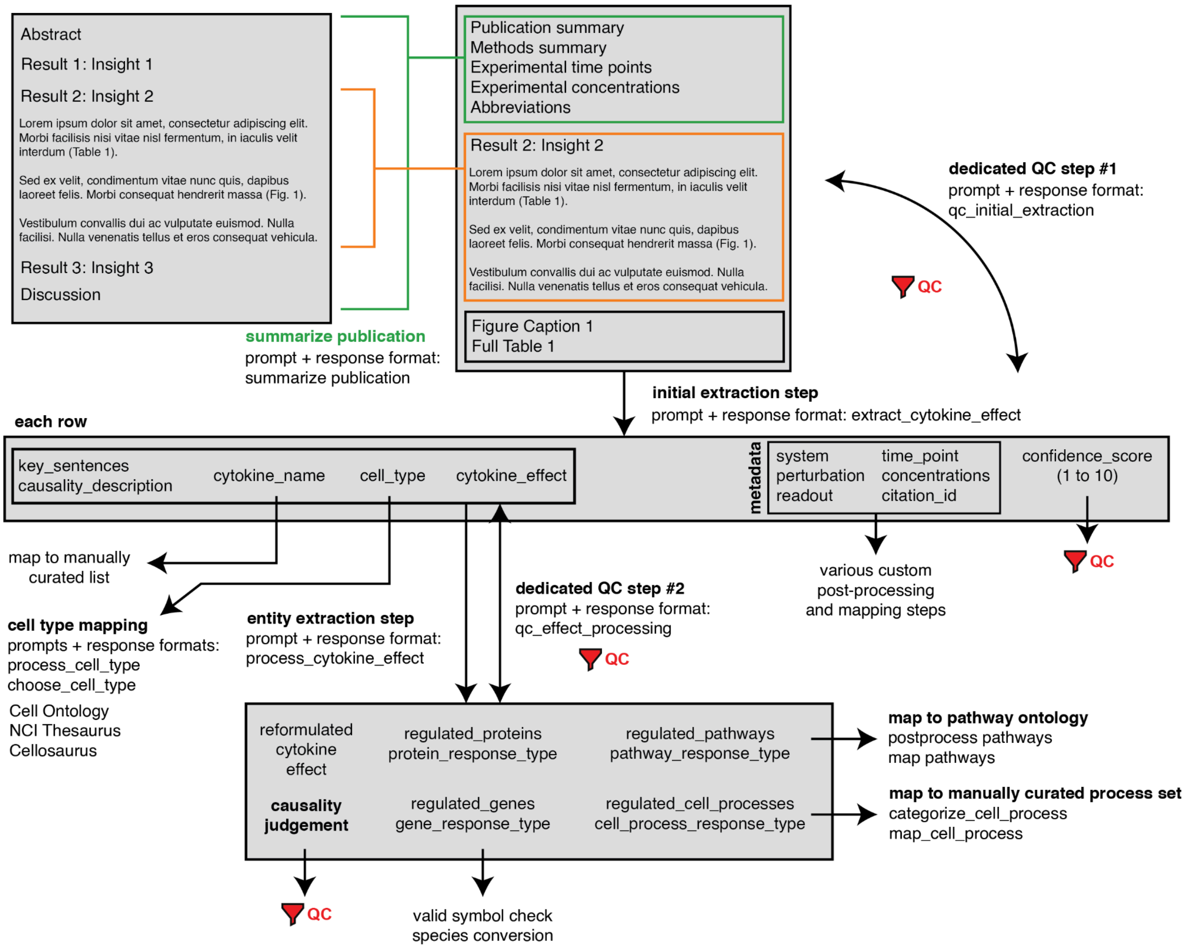


**Fig. S2. Schematic overview of the database construction workflow.** Major extraction, quality control, and standardization steps are shown along with their used input data and used prompts (see **Table S3** for details).

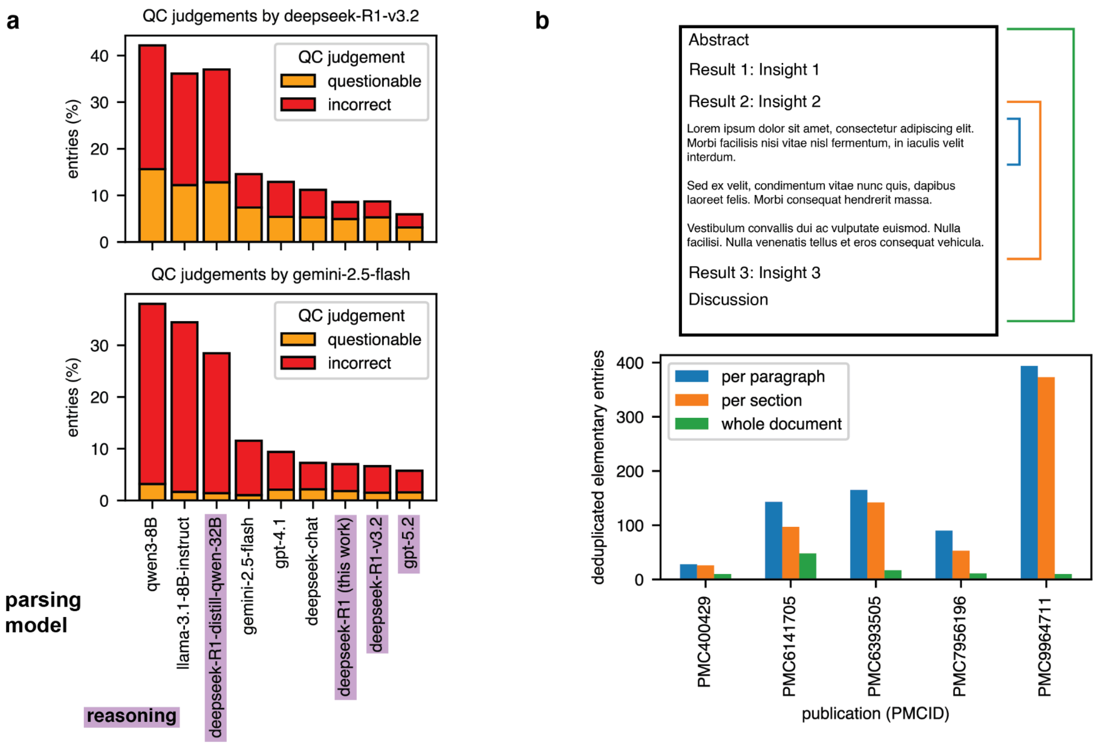


**Fig. S3. The choice of the parsing model and the publication chunking strategy. a,** Number of entries judged by two different LLMs (DeepSeek-R1-v3.2 or Gemini-2.5-Flash (medium reasoning)) as questionable or incorrect after the initial parsing step (prompt: qc_model_comparison). Models show little self-bias and have consistent rank ordering. Models that used reasoning in the parsing task are highlighted. Note that the derived error rate is not meant to represent an actual error rate relative to ground truth but to produce valid relative rankings between models. **b,** Number of unique triples per publication (deduplicated by a mixture of similarity-based grouped and LLM judgement, see Methods) for per paragraph, per section, or whole document chunking during the parsing process. While processing the whole document leads to severe under-extraction, per section chunking provides almost the same number of extracted statements while conserving greater context and requiring far fewer API calls. Per section chunking (with added context) is therefore chosen for this publication.

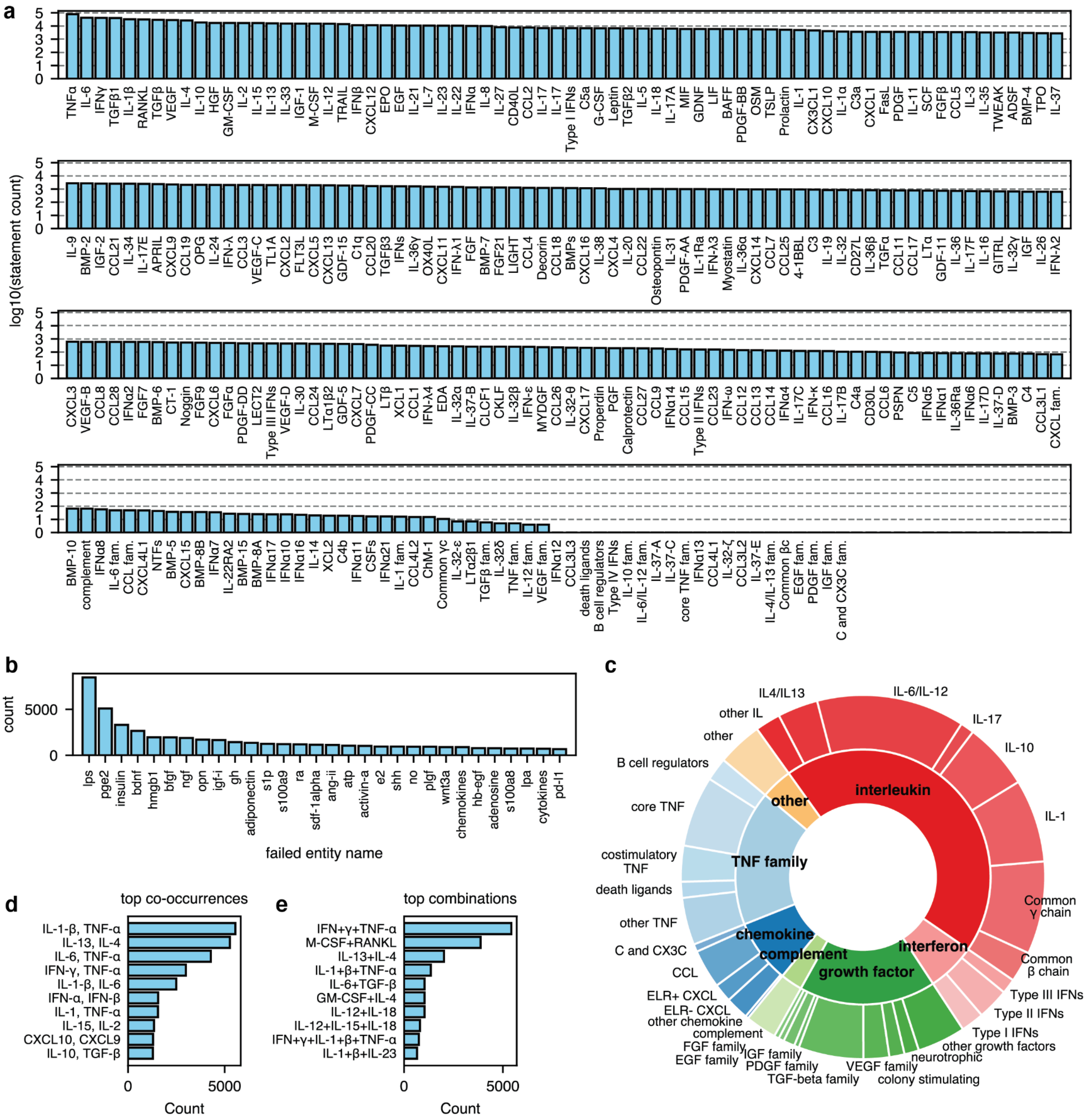


**Fig. S4. Statistics on extracted and mapped cytokines. a,** Count of triples per cytokine. **b,** Most common failed cytokine mappings. **c,** Distribution of extracted cytokines by family and subfamily. **d,** Most common co-occurring cytokines, i.e., those that are annotated as having the same effect for a particular triple. **e,** Top synergistic cytokines, i.e., those that are annotated as jointly producing an effect for a particular triple.

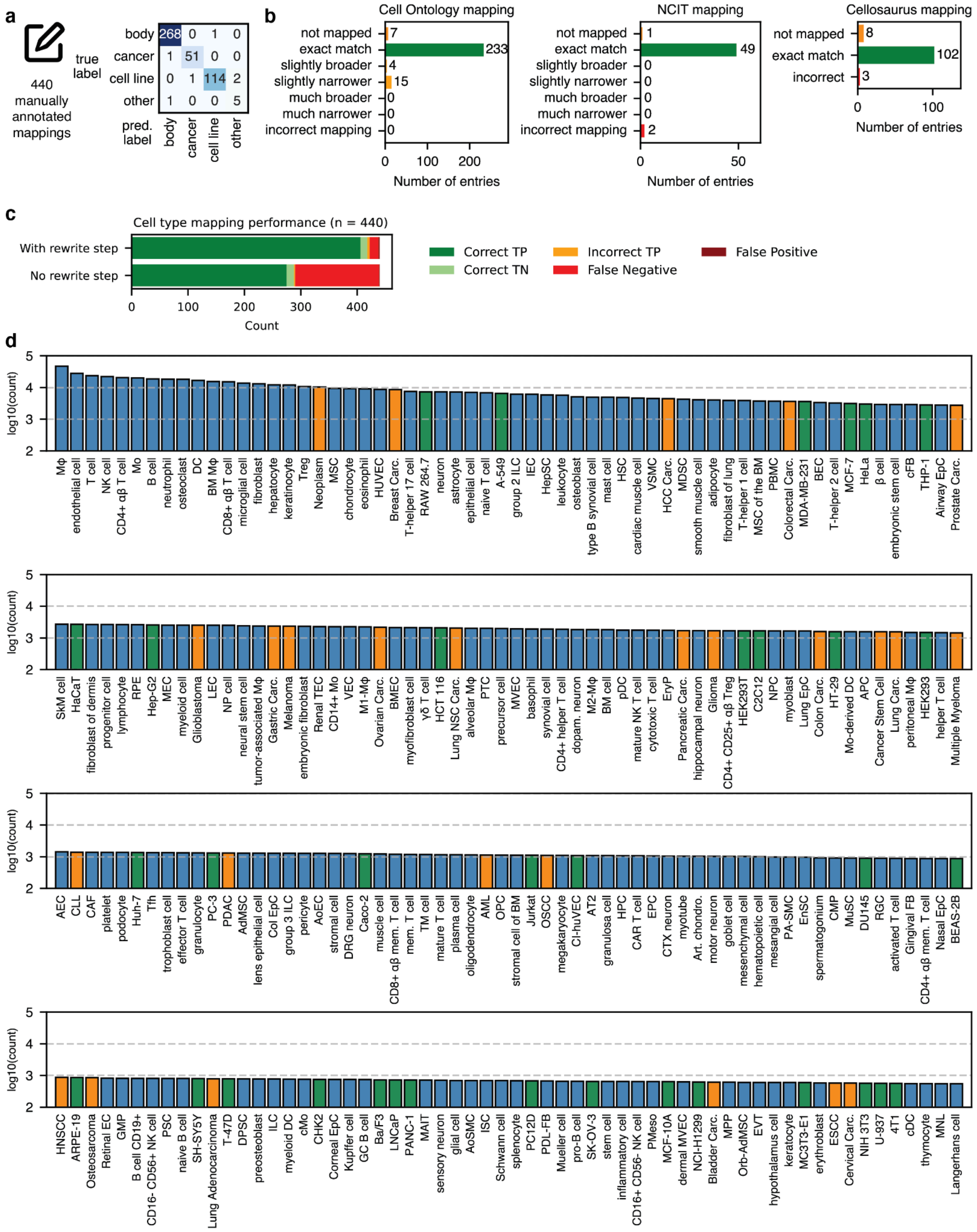


**Fig. S5. Statistics on mapped cell types. a,** Accuracy of mapping cell types to categories for a set of 440 manually annotated entries. **b,** Mapping accuracy to Cell Ontology, NCI Thesaurus, and Cellosaurus. For Cell Ontology and NCI Thesaurus, we distinguish slightly broader and slightly narrower (i.e., distance of at most 2 straight up or down the particular tree) from more distant or lateral off-target mapping. **c,** Comparison of cell type mapping performance on 440 manually annotated entries when applying the intermediate LLM-rewriting step ('With rewrite step') versus using the cell types from the paper directly ('Without rewrite'). True/False negatives refer to cell types where the ground truth annotation recognizes that it does not fit into our set of ontologies. **d,** Top mapped cell types by triple count.

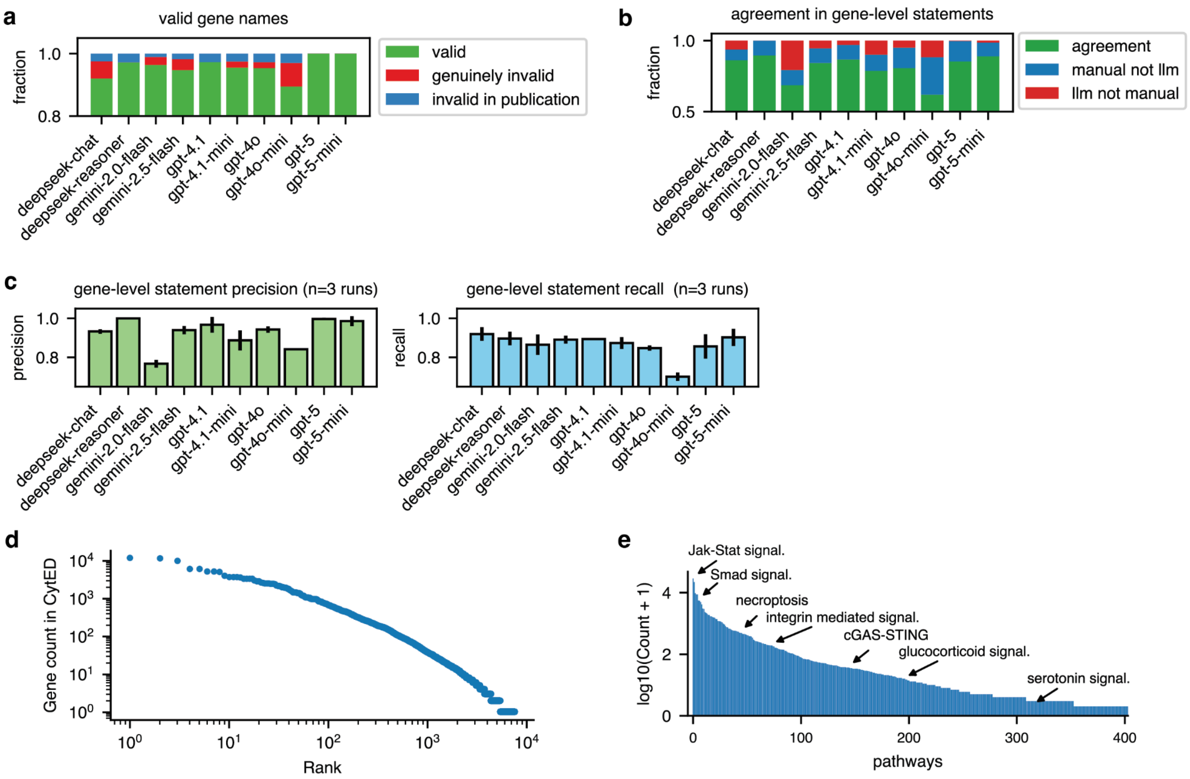


**Fig. S6. Evaluation of cytokine effect reformulation performance by different large language models. a,** Fraction of valid and invalid gene names in a manual set of 52 manually reformulated rows. Note that some gene names were already invalid in the original publication. **b,** Agreement in gene-level statements between manually and LLM-reformulated rows. The manual rows were explicitly constructed to contain all reasonable gene-level inferences, making additional statements likely errors. **c,** Precision (common statements / common statements + LLM-only statements) and recall (common statements / common statements + manual only statements) for different LLMs. **d.** Distribution of gene counts in CytED. **e.** Distribution of pathway counts in CytED after mapping.

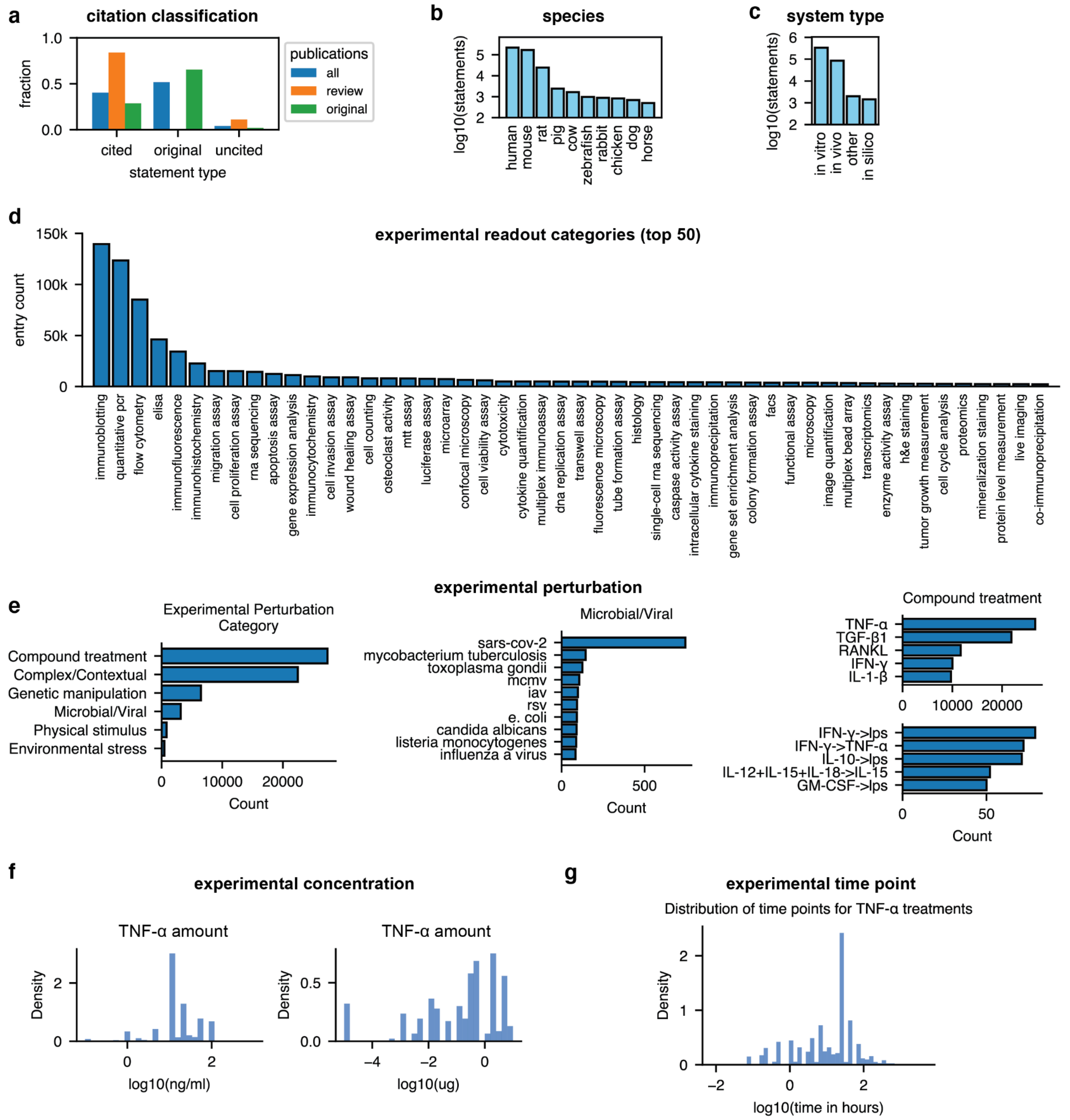


**Fig. S7. An overview over extracted metadata and its standardization. a.** Count of cited, original, and uncited (non-original) triples in all publications, reviews, and original publications. **b,** Count of statements by species. **c,** Count of statements by the experimental system type. **d,** Count of statements by the experimental readout category**. e,** Count of statements by the category of experimental perturbation (left), a particular viral challenge (center), and the most common compound treatments. The notation of, e.g., IL-12+IL-15+IL-18->IL-15 implies that the experiment initially used the first three simultaneously, then again IL-15 later. **f,** Distribution of experimental concentrations or amounts of TNFα across experiments. **g,** Distribution of experimental time points used for exogenous TNFα treatment experiments.

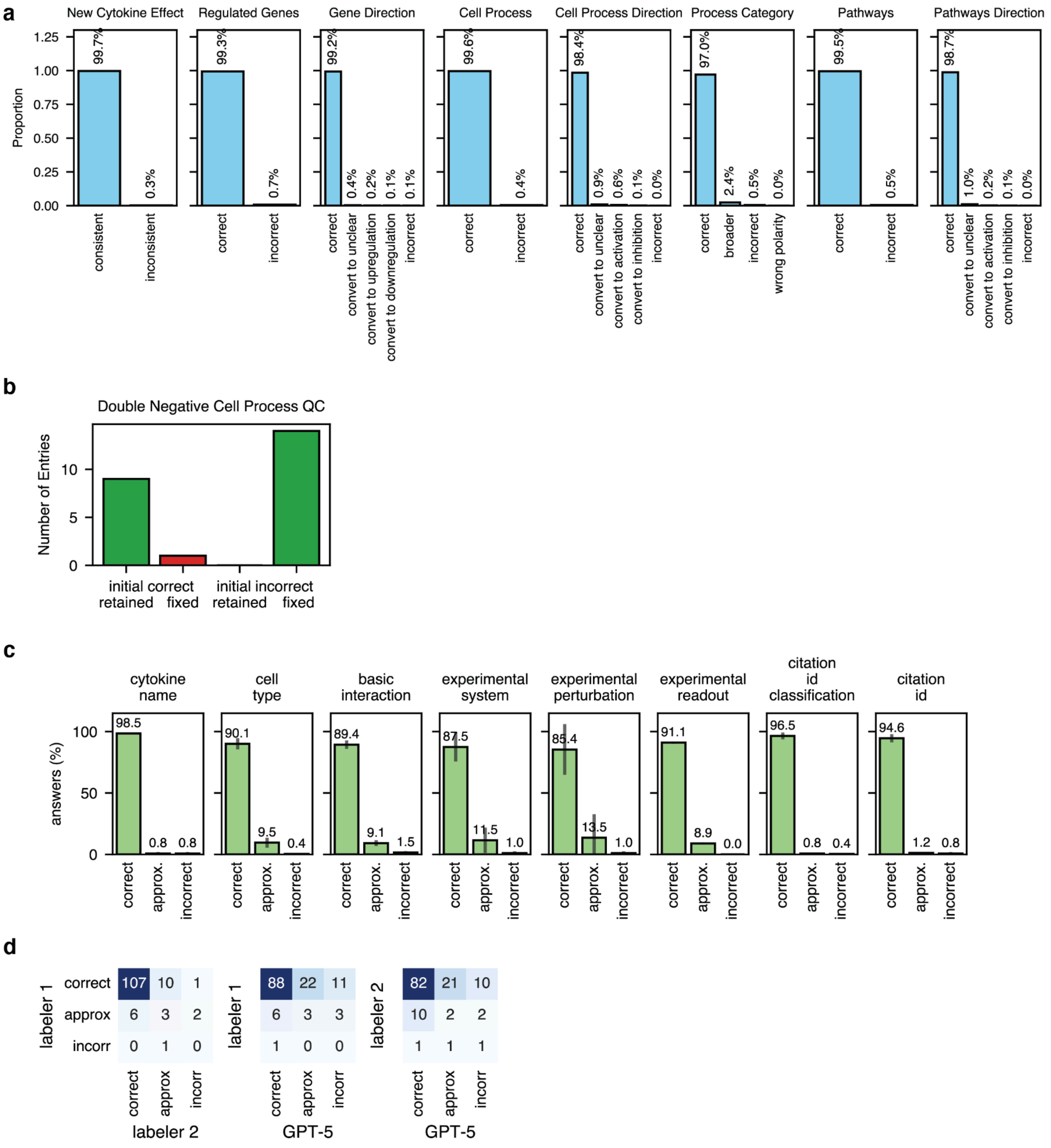


**Fig. S8. Additional quality control metrics. a,** Outputs of the quality control step for cytokine effect processing. **b,** Twenty manually labeled cases of double negatives in the cell effect column (e.g., 'inhibition of 'apoptosis inhibition'). Such double negatives had a particularly high likelihood of being incorrect. Almost all correct double negative entries are retained and all incorrect double negatives are removed by the reformulation QC. **c,** Distribution of manual labels in the quality control of 134 entries sampled to increase the baseline error rate. **d,** The concordance in the rating of the basic interaction described by entries is low both between labelers and between labelers and GPT-5. This indicates, to our mind, that few entries are obviously correct and entries labeled incorrect are largely edge cases.

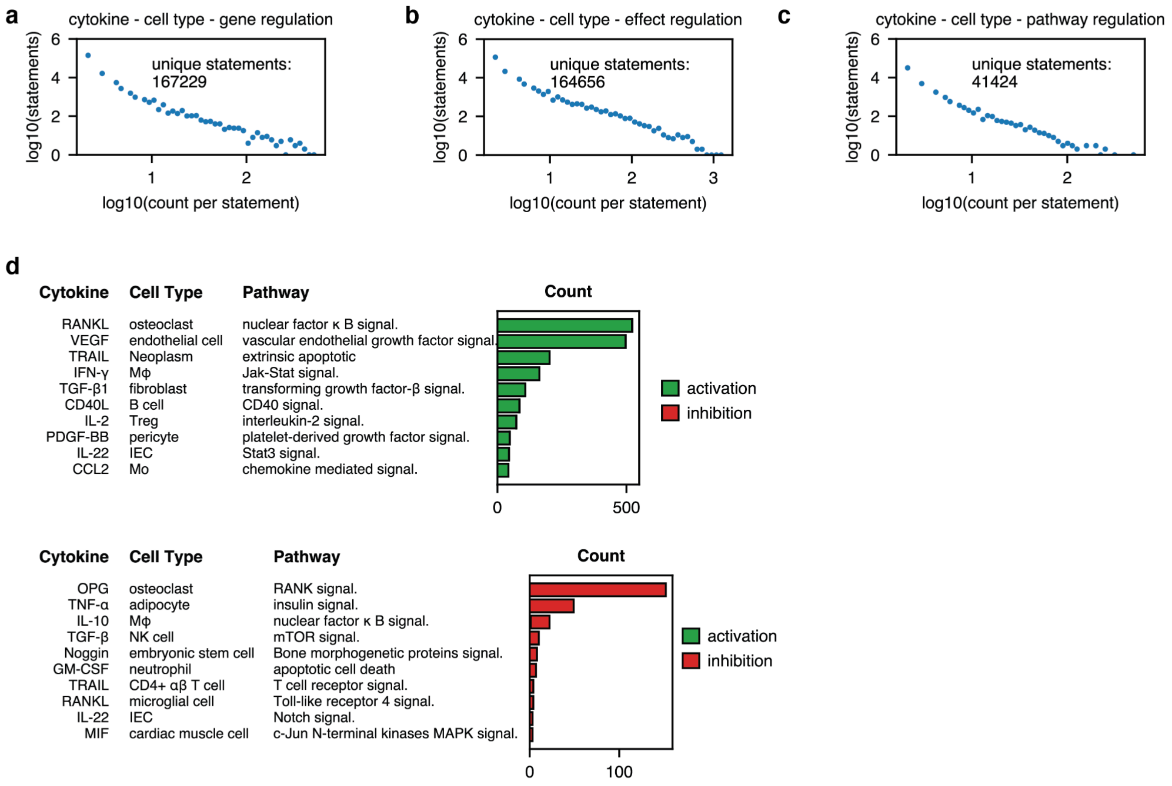


**Fig. S9. The distribution of literature support for cytokine effect statements. a, b, c,** Number of literature counts for individual (a) gene-, (b) cell effect- (c) pathway-level statements. **d.** Most commonly mentioned pathway activation and inhibition statements.

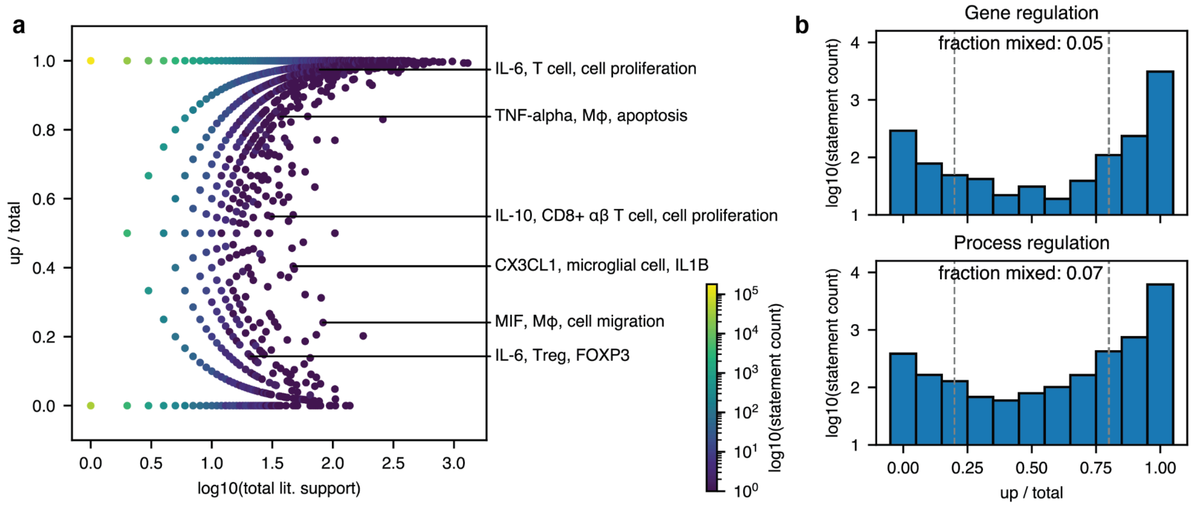


**Fig. S10. The context sensitivity of gene and cell effect statements. a,** Ratio of up-and down-regulation statements on both a gene- and cell effect-level and literature support. Some individual statements are highlighted. **b,** Distribution of the number of statements by the ratio of up- and downregulation for statements discussed in at least 6 publications. Statements with less than 80% agreement are classified as mixed.

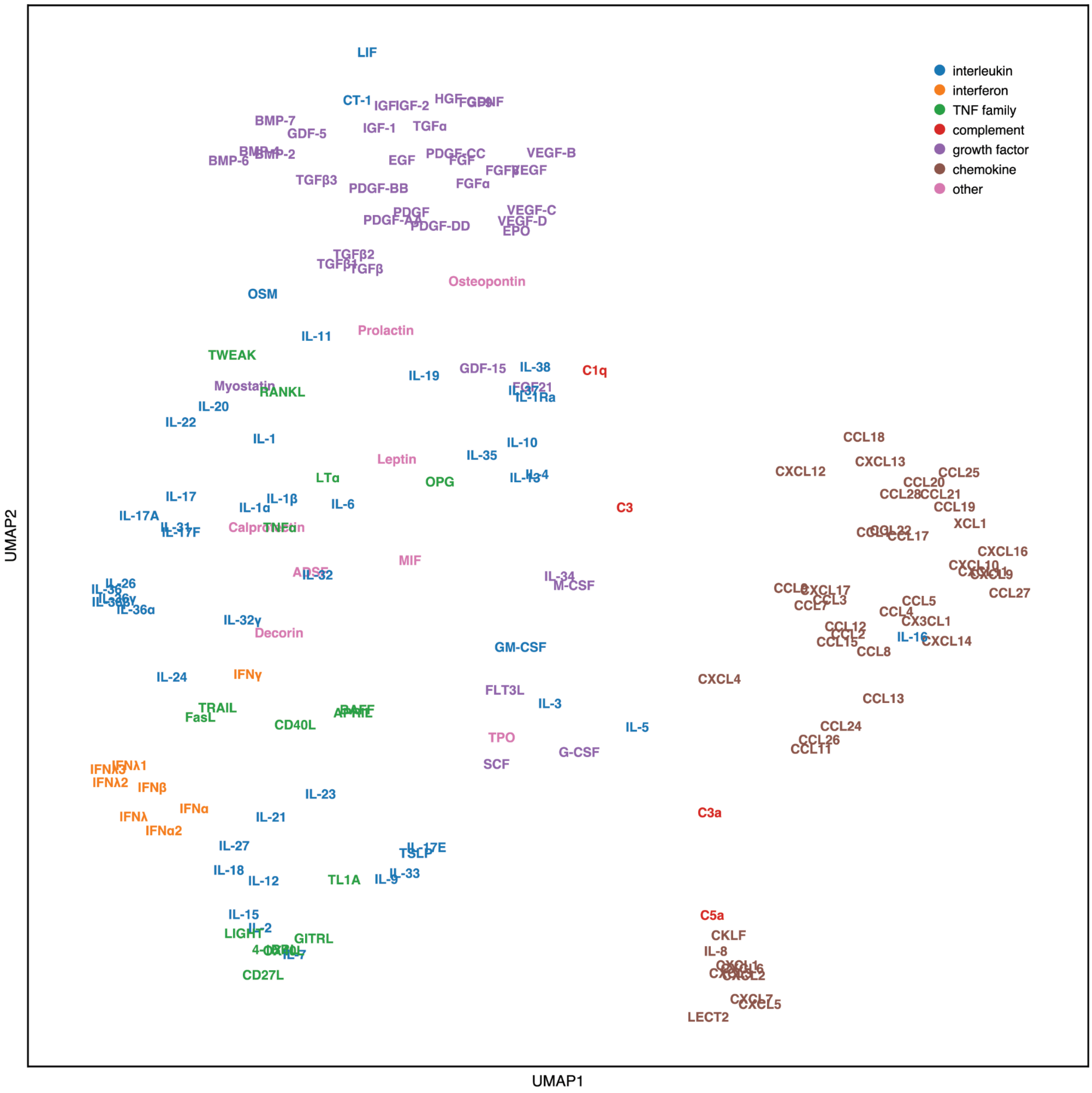


**Fig. S11. The distribution of cytokines by their effects in a two dimensional embedding.** UMAP of different cytokines for log10+1-transformed net literature counts of merged cell type-cell effect and cell type-gene relationships after truncated singular value decomposition.

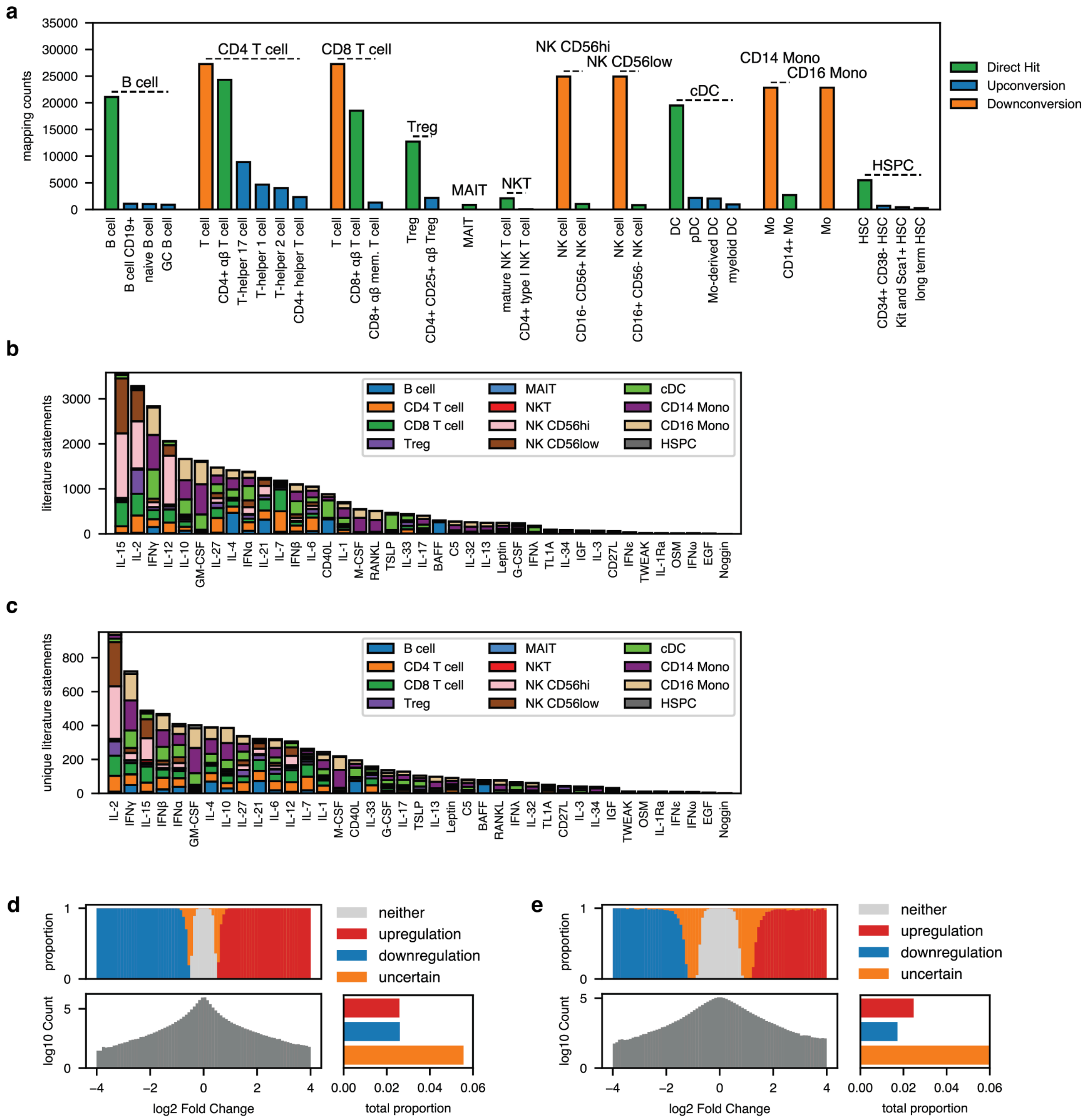


**Fig. S12. Integrating CytED with experimental data. a**, Overview over which cell types in CytED were either upconverted to a more general cell type or downconverted to a more specific cell type to match the particular cell types of the PBMC experimental dataset (indicated above the bars). **b,** Number of individual gene-level statements present both in CytED and in the differential expression data in the human PBMC dataset by both cell type and cytokine. **c,** Entries as in b but filtered to unique statements. **d.** Experimental results from a human PBMC screen are binarized based on abs(log2FC)>0.5 as indicating upregulation, downregulation, or no regulation of a specific gene in a cell type by a cytokine perturbation. Data points that are not confidently in any category are labeled uncertain and discarded. **e.** As in (d) but for an in vivo perturbation screen of mice. Binarization uses abs(log2FC)>1 due to higher noise levels in the data.

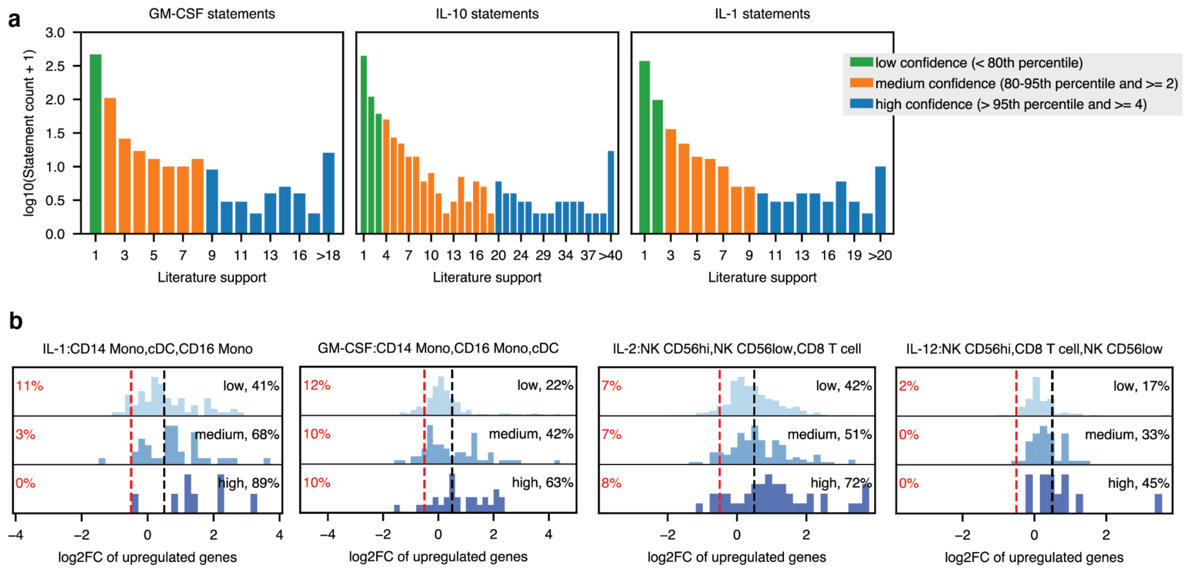


**Fig. S13. Assignment of confidence levels to statements and the impact of confidence levels on agreement with experimental results. a,** Number of statements by literature support and assigned confidence levels for GM-CSF, IL-10, and IL-1**. b**, Directional agreement (up- or downregulation) with the experimental data for different confidence levels for the most common target cell types for four selected cytokines.

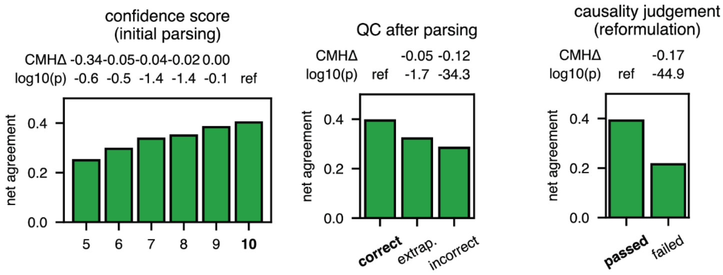


**Fig. S14. Quality control metrics inform the likelihood of experimental agreement.** Net agreement (same-opposite) between CytED and experimental data depending on the confidence score in the initial parsing, whether or not entries pass the post-parsing QC, and the causality judgement during reformulation. Statistical significance was determined via a generalized Cochran-Mantel-Haenszel (CMH) test stratified by cytokine identity relative to the reference group (ref.), with the weighted mean score difference (CMHΔ) as the effect size. Multiple testing was corrected using the Benjamini-Hochberg procedure.

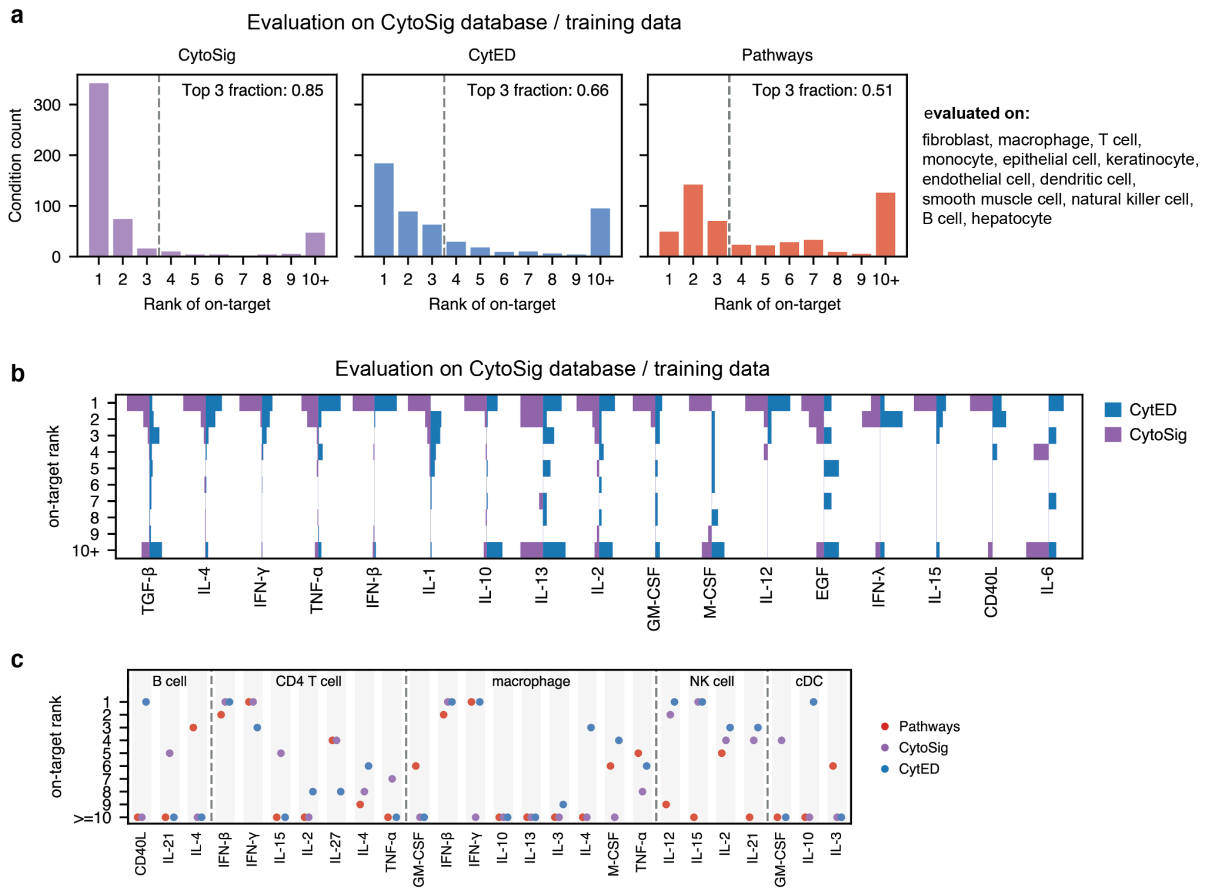


**Fig. S15. Additional evaluation results for cytokine signaling activity inference. a.** Distribution of on-target rankings for datasets in the CytoSig database/training data for the subset of cell types indicated on the right (n=507 datasets total). **b.** Distribution of rankings per cytokine for data as in (a). **c.** Ranking of the on-target cytokine for pathway-derived, CytoSig, and CytED-derived weighted gene sets in a subset of perturbation conditions for mouse in vivo data.

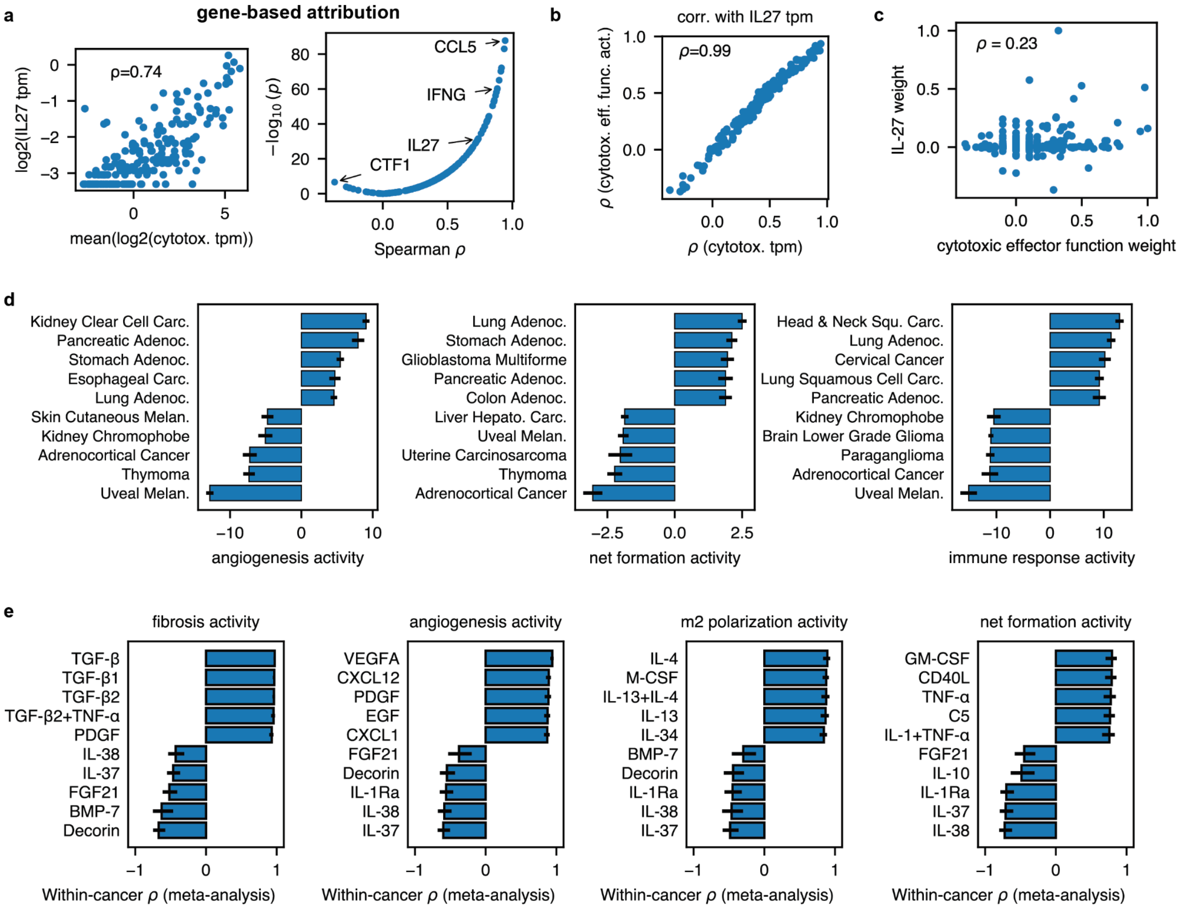


**Fig. S16. CytED infers process and cytokine activities in tumor samples. a.** Association between the mean of cytotoxic gene expression (*CD8A, GZMA, GZMB, GZMH, GZMK, PRF1, NKG7)* and *IL27* expression in melanoma samples from the TCGA dataset as well as the spearman correlation for the same set of cytotoxicity genes for various cytokine genes. **b.** Comparison of the correlations as in (a) derived from either the mean of cytotoxicity genes or inferred cytotoxic effector function process activities. **c.** IL-27 activity weights versus cytotoxic effector function weights per gene. **d.** Top and bottom five process activities across tumor types for three processes. Error bars indicate the 95% confidence interval across TCGA samples per cancer type. **e.** Top and bottom cytokine activity associations with four different processes TCGA. Bars show meta-analyzed within-cancer Spearman correlations between inferred cytokine activity and process activity across cancer types. Error bars indicate the standard deviation across cancer types.

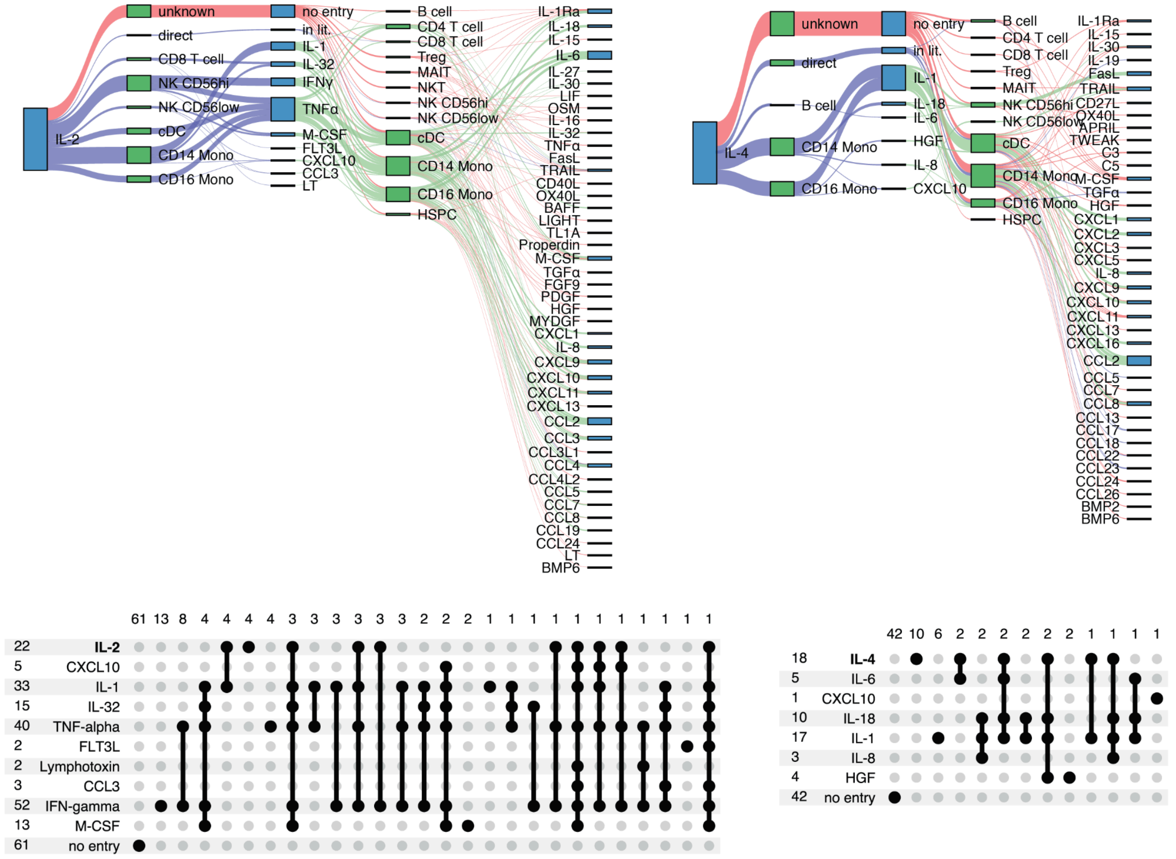


**Fig. S17. Additional examples of secondary effect inference.** Inference of primary and secondary effects of (left) IL-2 and (right) IL-4 based on CytED literature annotations. The inset shows how many connections are explained by the primary cytokine, secondary cytokines, or are not annotated in CytED for any of the cytokine perturbations.

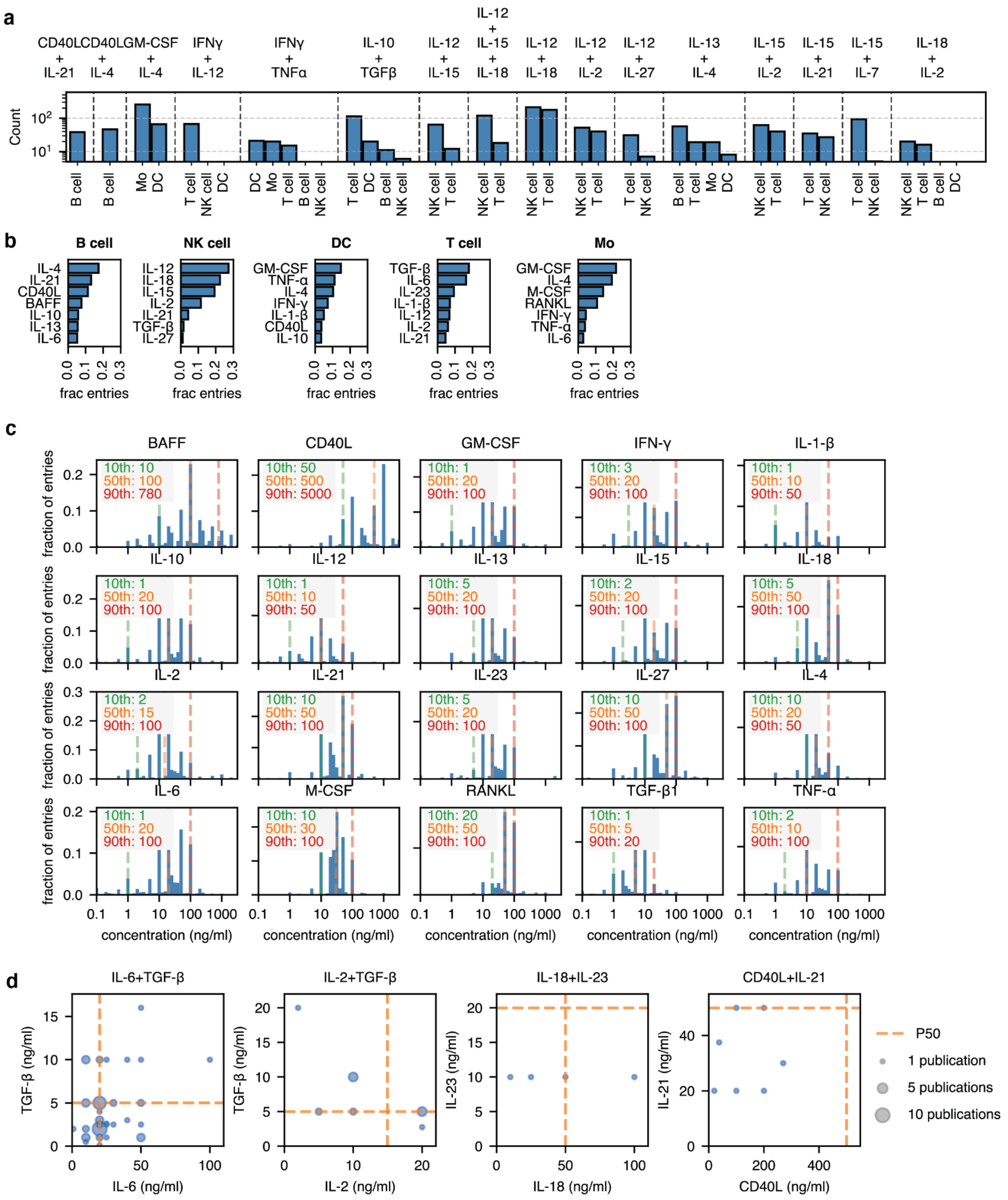


**Fig. S18. Picking core cytokines and cytokine combinations for a combinatorial PBMC screen. a,** Publication count in target cell types for common combinations. **b,** Most common individual cytokines that occur in cytokine combinations for PBMC cell types. **c,** Distribution of concentrations used in *in vitro* screens for each of the 20 most prominent combination cytokines. The 10th, 50th, and 90th percentile is shown as an inset on the top left of each panel. **d,** Most common concentrations used in *in vitro* screens for a representative set of cytokine combinations. The 50th percentile for individual cytokine perturbations is shown as a dashed line.

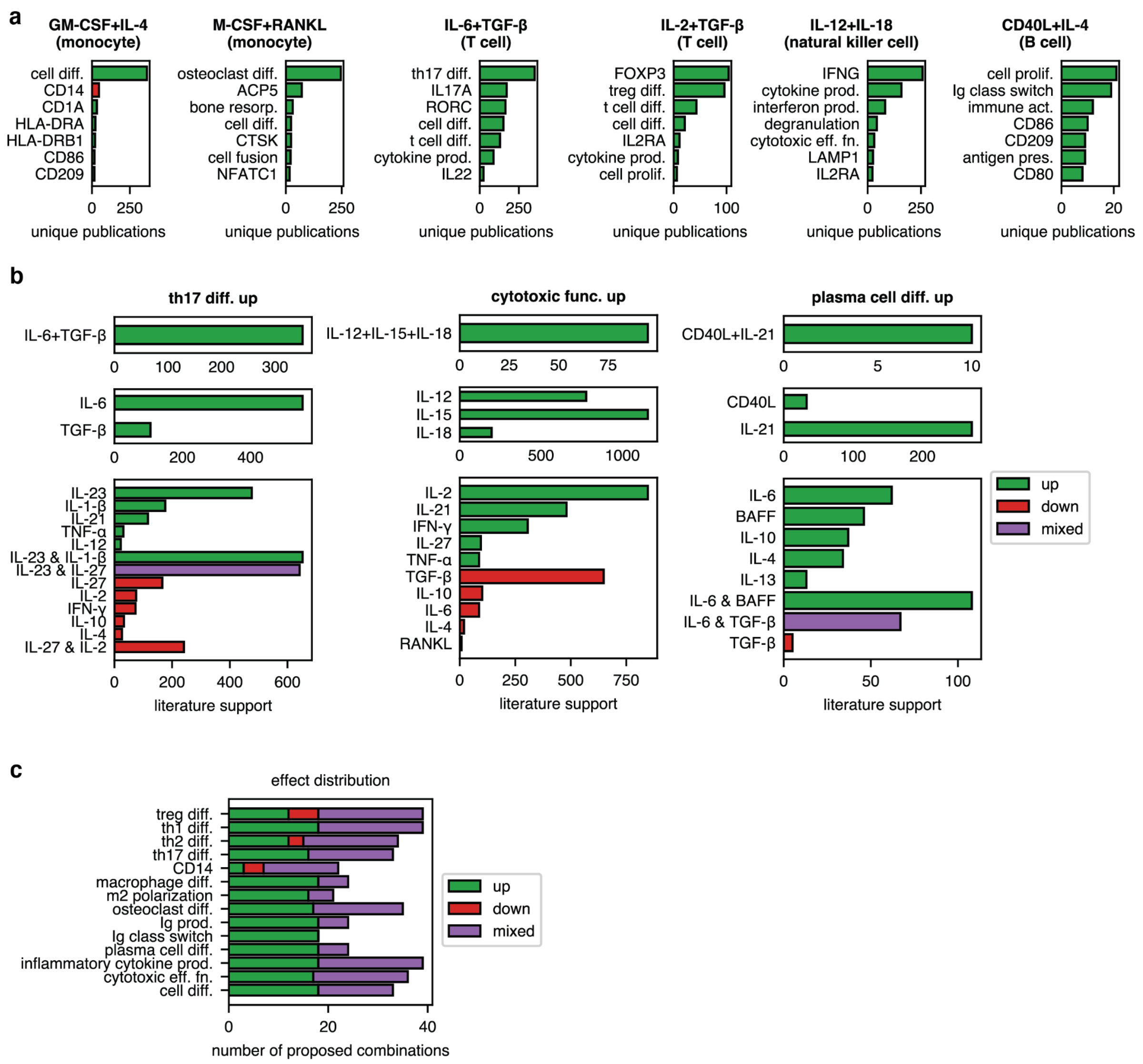


**Fig. S19. Identification of informative cytokine combinations by extending known interactions. a,** Representative cytokine combinations in specific cell types and the most frequently associated target processes or genes. **b,** Strategy for selecting novel combinations for a given target effect. The top panel shows the cytokine combination most commonly reported to regulate the indicated process or gene. The middle panel shows literature support for each individual cytokine affecting the same target. The bottom panel shows additional cytokines predicted to form novel combinations with the original set based on shared regulation of the target process/gene. **c.** Number of picked novel cytokine combinations for each target process or gene, stratified by whether the constituent cytokines collectively activate, inhibit, or exhibit mixed effects on the target process or gene.
